# Myofibroblastic CAFs arising from bone-resident osteoblast precursors retain an osteolineage signature and support breast cancer progression via Osterix-mediated signaling

**DOI:** 10.1101/2025.09.18.677154

**Authors:** Giulia Furesi, Carisa Zeng, Emily M. Eul, Deborah J. Veis, Jennifer Ye, Vasilios A. Morikis, Taylor Malachowski, Darya Khantakova, Alina Ulezko Antonova, Marco Colonna, Gregory D. Longmore, Sheila A. Stewart, Maxim N. Artyomov, Roberta Faccio

## Abstract

Cancer-associated fibroblasts (CAFs) are a major component of the breast cancer (BC) microenvironment, involved in tumor progression and resistance to therapy. Despite the recent identification of multiple CAF subtypes with unique functions, it remains unclear whether each subtype arises from a distinct precursor or a shared common progenitor. Here, we identified a unique subpopulation of myofibroblast CAFs (myCAFs) arising from committed Osterix (Osx)+ osteoblast progenitors in the bone, recruited to primary tumors in both murine BC models and BC patients. Osx^+^myCAFs exhibit strong protumorigenic features and retain osteoblastic gene expression, which distinguishes them from Osx^neg^CAF subsets. Osx drives the expression of extracellular matrix remodeling genes and promotes tumor growth via the secretion of MMP13, a key Osx target gene. Finally, we find that increased Osx^+^myCAFs and a stromal osteolineage gene signature correlate with poor therapeutic response and reduced BC patient survival.

## Introduction

Breast cancer (BC) remains the most frequently diagnosed cancer in women, accounting for approximately 30% of new cases each year in the United States. While genetic and molecular alterations within cancer cells are key drivers of tumorigenesis (1, 2), the tumor microenvironment (TME) plays an equally crucial role in shaping tumor behavior and response to therapy (3, 4). Cancer-associated fibroblasts (CAFs) are a heterogeneous group of stromal cells within tumors (5), best known for their roles in depositing and remodeling a collagen-rich extracellular matrix (ECM). Elevated ECM stiffness significantly impacts key biological processes involved in tumor progression, including enhancing tumor cell proliferation and migration, creating a barrier to immune infiltration, and inducing resistance to therapy (6). CAFs can also secrete chemokines and cytokines to modulate anti-tumor immune responses and produce factors that directly promote tumor cell proliferation and survival (7, 8). The presence of CAFs is associated with poor clinical outcomes in multiple cancer types, including BC, making them an attractive therapeutic target (9, 10). Nonetheless, their phenotypic diversity and resemblance to normal fibroblasts pose significant challenges to designing effective CAF-targeted treatments.

Like normal fibroblasts, CAFs express the stromal markers alpha-smooth muscle actin (α-SMA), fibroblast activation protein alpha (FAP-α), platelet-derived growth factor receptors (PDGFRβ and PDGFRα), and fibroblast-specific protein 1 (S100A4/FSP1). However, unlike normal breast fibroblasts, CAFs have specialized functions within the TME (11). Previous studies have utilized scRNA-seq analysis to classify CAFs into distinct subsets based on gene signatures and functions. These include myofibroblastic (myCAFs) and matrix-producing CAFs (mCAFs), involved in ECM production and remodeling (12, 13); inflammatory (iCAFs), antigen-presenting (apCAFs), and senescent CAFs (senCAF), linked to immune responses (14–17); vascular CAFs (vCAFs), which support vascular growth (13, 18); and additional specialized subsets, among others (13, 18). Yet, it is unclear whether these specialized CAF subsets originate from distinct progenitors or from a common precursor that adapts to local cues to acquire specific phenotypic traits.

Evidence supporting both a single universal progenitor and multiple cellular origins exists. Studies by Buechler et al. suggest that tissue-resident fibroblasts act as conserved progenitors, capable of transitioning into multiple CAF subtypes in response to tumor-derived signals due to their inherent plasticity (19). However, other studies indicated that CAFs can originate from bone marrow mesenchymal stromal cells (BMSC), adipocytes, pericytes, epithelial and/or endothelial cells, and even cancer stem cells (20–25). This variability might be cancer-type and/or cancer-stage specific (10), but whether the origin of CAFs dictates their phenotype/functionality in the TME needs further investigation.

Osteoblasts are bone-resident populations derived from BMSC, specialized in bone formation (26). Given the functional similarities between osteoblasts and CAFs, particularly related to ECM production and remodeling, we hypothesized that committed osteoblast progenitors (osteolineage cells) located in the bone could give rise to matrix-producing CAFs in the TME. Osterix (Osx) is a transcription factor expressed by osteolineage cells, required for osteoblast differentiation and bone formation (27). With the use of Osx;Cre lineage tracing mouse models, the presence of a few Osx^+^ cells and/or their progeny has been detected outside of the bone, including in solid tumors (28, 29). Surprisingly, these initial observations indicated that over 90% of Osx-lineage derived cells in the TME comprised CD45^+^ immune populations, originated from rare Osx^+^ hematopoietic stem cells that lose Osx expression upon lineage allocation. Whether Osx was expressed and exerted any pro-tumorigenic function in the CD45^neg^ stromal populations residing in the TME was not pursued at the time. In this study, we identified a subset of myCAFs derived from Osx^+^ osteoblast precursors, retaining Osx expression and with superior pro-tumorigenic function compared to Osx^neg^CAFs. Osx controls the expression of MMP13, and its targeting selectively reduces growth of tumors supported by Osx^+^myCAFs. Notably, the presence of Osx^+^myCAFs in BC tissues correlates with a poor response to therapy and overall reduced survival in BC patients.

## Results

### Osteoblast progenitors migrate from the bone marrow to the TME to enhance tumor growth

To determine if tumor-infiltrating CAFs involved in the ECM deposition originate from committed Osx^+^ osteoblast progenitors residing in the bone, we used the doxycycline (doxy)-repressible OsxCre;TdT lineage tracing model (30). Mice were fed a doxy-diet until weaning to activate the Cre in committed osteoblast progenitors in bone marrow (BM), mature osteoblasts, and osteocytes. 8-week-old OsxCre;TdT female mice were then orthotopically inoculated with the luminal B, ER⁺/PR⁺, hormone-resistant PyMT cell line, and the presence of CD45^neg^TdT^+^ cells (designated as TdT^+^OsteoLin cells) in circulation and at tumor site was analyzed two weeks later. We detected a small pool of TdT^+^OsteoLin cells in blood of non-tumor bearing mice (NTB), representing ∼0.2% of total blood cells, and observed a five-fold increase following tumor inoculation (Fig. 1A, Suppl. Fig. 1A). When we examined the primary tumor, we observed that TdT⁺OsteoLin cells, comprising 0.8–1.2% of the total tumor mass (Suppl. Fig. 1B), colocalized with PDGFRβ, a pan-mesenchymal and CAF marker (Fig. 1B). Flow cytometry analysis further confirmed that over 80% of TdT⁺OsteoLin cells were PDGFRβ⁺ (Fig. 1C, Suppl. Fig. 1C). Interestingly, PDGFRβ^+^;TdT^+^OsteoLin cells comprised only ∼20% of PDGFRβ^+^ stromal cells within the tumor mass (Fig. 1D), suggesting they represent a distinct subset of tumor-infiltrating mesenchymal cells.

**Figure 1.**
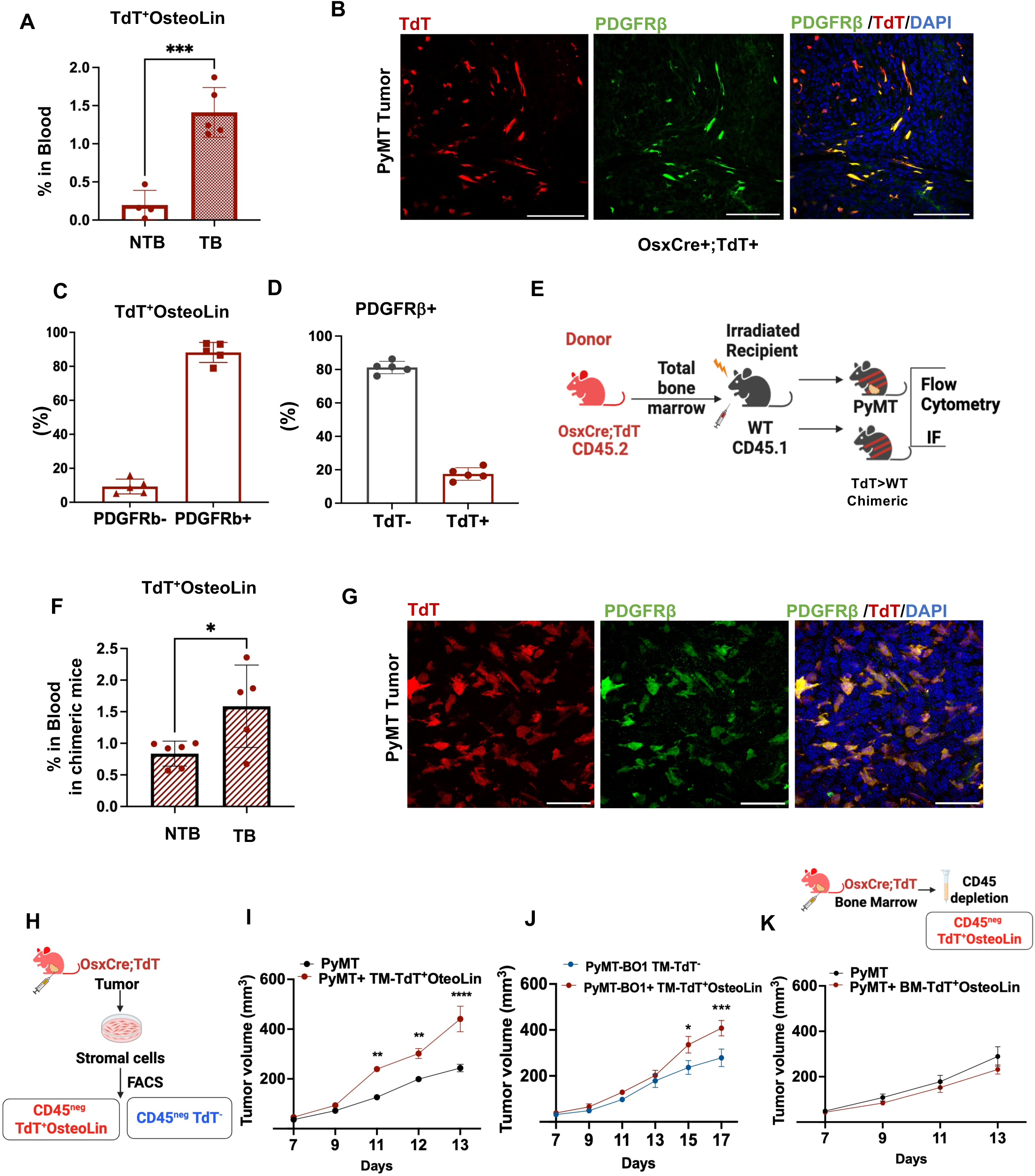
Osteoblast progenitors migrate from the bone marrow to the TME to enhance tumor growth. **A,** Flow cytometry analyses showing the percentage of TdT^+^OsteoLin in circulation of no-tumor (NTB) and tumor-bearing OsxCre;TdT mice (TB) orthotopically inoculated with PyMT cells. **B,** Representative IF images of PyMT tumors showing TdT^+^OsteoLin (red), PDGFRβ (green), and DAPI (blue). Bar=50μM. **C,** Flow cytometry analysis showing the percentage of TdT^+^OsteoLin cells expressing PDGFRβ in the TME of PyMT-BO1-GFP^+^ orthotopic tumors. **D,** Flow cytometry analyses showing the percentage of PDGFRβ^+^ expressing TdT in the TME of PyMT-BO1-GFP^+^ orthotopic tumors. **E-F,** Schematic representation showing transplantation of bone marrow cells from OsxCre;TdT mice into irradiated CD45.1 recipient WT animals to generate chimeric TdT>WT mice (F). Chimeric mice were orthotopically injected 6 weeks later with PyMT cells in the mammary fat pad and subjected to IF and flow cytometry to detect donor-derived TdT^+^OsteoLin^+^ cells. (J) Flow cytometry analysis showing the percentage of CD45^neg^TdT^+^ in blood from chimeric tumor-bearing and no tumor-bearing mice in (I). **G,** Representative IF z-stack images of PyMT tumors in chimeric mice stained for PDGFRβ (green) and DAPI (blue) with TdT^+^ cells displayed in red. Bar=50μM. **H,** Model showing isolation of tumor-derived (TM) stromal cells by plating dissociated orthotopic tumors from OsxCre;TdT mice on tissue culture dishes for 30min, followed by cell sorting of the adherent GFP^neg^CD45^neg^TdT^+^OsteoLin and GFP^neg^CD45^neg^TdT^neg^ populations. **I-J,** Tumor growth determined by caliper measurements in WT mice inoculated with indicated tumor cells alone or co-injected with CD45^neg^TdT^+^OsteoLin cells at 1:1 stroma:tumor ratio (M) or GFP^neg^CD45^neg^TdT^+^OsteoLin and GFP^neg^CD45^neg^TdT^neg^ at 1:2 stroma:tumor ratio (N) (n=6/group). **K,** Model illustrating the isolation of BM-derived stromal cells from OsxCre;TdT mice with orthotopic PyMT tumors, after exclusion of CD45⁺ cells by magnetic beads, followed by sorting of the BM CD45^neg^TdT^+^OsteoLin cells (top). Tumor growth was determined by caliper measurement in WT mice injected with tumor cells alone or co-injected with BM-derived TdT^+^OsteoLin cells (n=6/group). Experiments were conducted in at least three independent replicates. Results are shown as mean +/- SD (A, C, D, F) or SEM (I, J, K). Unpaired Student T-test was performed to determine significance in A and F, Two-way ANOVA followed by Turkeýs multiple-comparison test (I, J, K) was used to assess significance (* P≤0.05, ***P≤0.001, ****P≤0.0001).

To demonstrate that the tumor-infiltrating TdT^+^OsteoLin cells are mobilized from the BM, we transplanted BM from OsxCre;TdT^+^ mice (CD45.2) into naïve, lethally irradiated wild-type (WT) recipients (CD45.1) (Fig. 1E). In contrast to classical bone marrow transplantation (BMT) used for hematopoietic purposes, we transplanted five times the standard amount of BM cells to ensure transfer of mesenchymal populations. Chimerism was confirmed by the presence of over 80% CD45.2^+^ immune cells in circulation starting three weeks post-transplant (Suppl. Fig. 1D, 1E). The chimeric mice were orthotopically injected with PyMT tumor cells at six weeks post-BMT, and the presence of donor-derived TdT^+^OsteoLin cells was assessed two weeks later in the blood and tumor mass. We observed a 1.9-fold increase in the number of TdT^+^OsteoLin cells in the blood of chimeric tumor-bearing mice versus chimeric no-tumor controls (Fig. 1F). Importantly, we detected PDGFRβ^+^; TdT^+^OsteoLin cells in the tumor mass (Fig. 1G, Suppl. Fig. 1F), confirming that tumor-infiltrating mesenchymal TdT^+^OsteoLin cells originate from an Osx^+^ osteoprogenitor population in the BM.

Next, to investigate the impact of TdT^+^OsteoLin cells on tumor growth, we isolated CAF populations from orthotopic PyMT or GFP^+^PyMT-BO1 tumors in OsxCre;TdT mice by plating the dissociated tumor mass on tissue culture dishes for 30min and removing the cells in suspension, mainly composed by tumor cells and immune populations, as previously shown (7) (Fig. 1H). Adherent TdT^+^OsteoLin cells were sorted based on TdT expression and re-injected with PyMT tumor cells at 1:1 stroma-tumor ratio into age-matched WT recipient animals. Notably, tumor cells co-injected with TdT^+^OsteoLin cells grew faster than PyMT cells alone (Fig. 1I). To assess whether TdT^+^OsteoLin cells share functional similarities with other tumor-infiltrating mesenchymal populations, we co-injected PyMT-BO1 tumor cells with GFP^neg^CD45^neg^TdT^+^OsteoLin or GFP^neg^CD45^neg^TdT^neg^ cells, sorted from PyMT-BO1-GFP orthotopic tumors, into new recipient WT animals (Suppl. Fig. 1G). To our surprise, TdT^+^OsteoLin cells showed superior pro-tumorigenic effects compared to their TdT^neg^ counterpart (Fig. 1J). In contrast, co-injection of PyMT tumor cells with TdT^+^OsteoLin cells isolated from the BM of mice bearing orthotopic PyMT tumors into WT recipient mice failed to increase tumor growth compared to PyMT alone (Fig. 1K). These results indicate that tumor-infiltrating mesenchymal TdT^+^OsteoLin cells derive from a BM resident Osx^+^ osteoblast progenitor that acquires significant tumor-supporting abilities after reaching the TME.

### Tumor-infiltrating TdT^+^OsteoLin cells display a myCAF-like phenotype

To assess the identity of tumor-infiltrating TdT^+^OsteoLin cells, we sorted GFP^neg^CD45^neg^Ter119^neg^CD31^neg^TdT^+^ or TdT^neg^ fractions from PyMT-BO1-GFP^+^ orthotopic tumors in OsxCre;TdT mice (Fig. 2A) and performed single-cell RNA sequencing (scRNA-seq). Because of the technical limitations that may arise from having a lower percentage of TdT^+^ compared to TdT^neg^ cells, we performed 10X 3’ scRNA-seq on sorted CD45^neg^ populations (including TdT^+^ and TdT^neg^ fractions) and CD45^neg^TdT^+^ cells, pulled from different cohorts of mice (Fig. 2A). Both datasets were merged before analysis and visualized using the Uniform Manifold Approximation and Projection (UMAP) algorithm. Cluster identities were manually annotated based on established expression marker profiles (Suppl. Fig. 2). We detected three major clusters of stromal cells (Fig. 2B): CAFs (cluster 0: *Pdgfrβ, Pdgfrα, Fap, Acta2),* Pericyte/vCAF (cluster 5: *Myh11, Vtn, Mcam)*, and endothelial cells (cluster 3: *Tie1, Icam2, Flt4, Pcam1*). Despite sorting for GFP^neg^ and CD45^neg^ stromal populations, we also identified small clusters of epithelial cells (cluster 1: *Bnc1, Krt8, Krt18, Msln*), macrophages (cluster 2 *Ptprc, Adgre1, Fcgr1, C1qb*), and dendritic cells (DC) (cluster 8: *Ptprc, Flt3, Zbtb46, Batf3*) (Fig. 2B and Suppl. Fig. 2).

**Figure 2.**
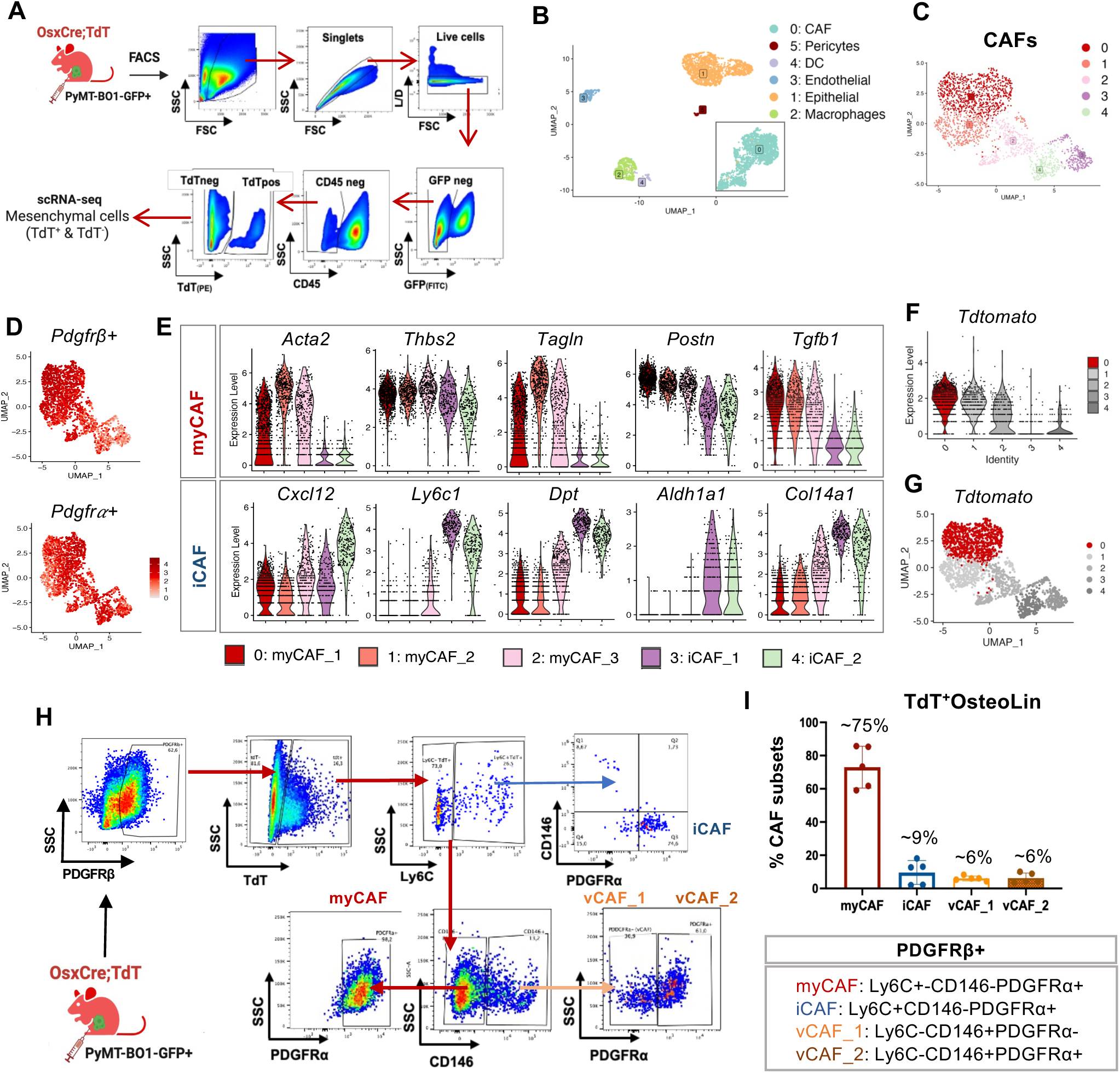
Tumor-infiltrating TdT^+^OsteoLin cells represent a distinct subset of myCAFs. **A**, Schematic representation of tumor-infiltrating TdT^+^ and TdT^neg^ stromal populations sorted from PyMT tumors eleven days post orthotopic inoculation in OsxCre;TdT mice and used from scRNAseq. n=4 mice were pooled for sorting. **B,** UMAP of the murine scRNA-seq dataset showing indicated clusters. **C,** UMAP showing the reclustering of CAF subpopulations. **D,** Feature plot showing the expression of *Pdgfrβ* and *Pdgfrα* in CAFs. **E,** Violin plots of indicated CAF subset markers. **F-G,** Violin plot UMAP (F) and UMAP (G) showing the murine myCAF subset with the highest TdT expression (red). **H-I,** Gating strategy for profiling TdT^+^OsteoLin CAF subpopulations via flow cytometry from GFP^+^PyMT-BO1 tumors in OsxCre;TdT+ mice (H). Gating shown after exclusion of GFP^+^, CD45^+^, Ter119^+^, CD31^+^ cells. (I), Distribution of CAF subsets expressed as a percentage of PDGFRβ^+^;TdT^+^OsteoLin cells. CAF markers used for analysis indicated in inset.

Given PDGFRβ expression in TdT^+^OsteoLin cells (Fig. 1C), we focused our analyses on the CAF cluster (cluster 0). UMAP dimensionality reduction and graph-based clustering separated five distinct subpopulations of CAFs (Fig. 2C), all of which were positive for PDGFRβ and PDGFRα, albeit to a different degree (Fig. 2D). By using additional established CAF markers detected in BC patients and the spontaneous PyMT-MMTV BC model (16, 31), we identified 3 major sub-clusters of myCAFs (myCAF1-3), and two of iCAFs (iCAF1-2). MyCAFs were marked by the expression of *Acta2*, *Thbs2, Tagln, Postpn,* and *Tgfβ1* and the iCAFs by the expression of *Cxcl12, Ly6C1, Aldh1a, Dept, and Col14a1* (Fig. 2E). Strikingly, TdT expression was mainly detected in the myCAF1 subset (Fig. 2F-G).

To validate this finding, we performed flow cytometry to assess the CAF composition in orthotopic PyMT-BO1-GFP^+^ tumors from OsxCre;TdT mice. PDGFRβ^+^ cells were selected after excluding tumor cells (GFP^+^), red blood cells (Ter119^+^), immune cells (CD45^+^), and endothelial cells (CD31^+^). Next, we gated on TdT+ cells and evaluated the surface expression of markers identified in myCAFs (PDGFRβ^+^, Ly6C^neg^, CD146^neg^, PDGFRα^+^), iCAFs (PDGFRβ^+^, Ly6C^+^, CD146^neg^, PDGFRα^+^), and vCAFs (PDGFRβ^+^, Ly6C^neg^, CD146^+^, PDGFRα^+^) (Fig. 2H). The majority of TdT^+^ cells (75%) clustered with myCAFs, with only a small percentage (<10%), expressing iCAF and vCAF markers (Fig. 2I). Altogether, these results indicate that tumor-infiltrating TdT^+^OsteoLin cells represent a subset of myCAFs derived from a BM Osx^+^ osteoblast progenitor.

### Osx^+^myCAFs are found in murine breast tumors and retain an osteoblastic signature

Given that Osx/Sp7 is a transcription factor whose detection is challenging by scRNA-seq, we next evaluated whether TdT^+^OsteoLin-derived myCAFs retain Osx expression. GFP^neg^;CD45^neg^;TdT^+^OsteoLin tumor-infiltrating cells were sorted from GFP⁺PyMT-BO1^+^ tumors in OsxCre;TdT mice and Osx expression was analyzed by IF. Nuclear localization of Osx was only observed in the TdT^+^OsteoLin cells, consistent with its role as an active transcription factor (Fig. 3A), but not in the GFP⁺ tumor cells (Suppl. Fig. 3). Based on this observation, we designated the tumor-infiltrating TdT^+^OsteoLin cells as Osx^+^myCAFs, a term we will use from here on.

**Figure 3.**
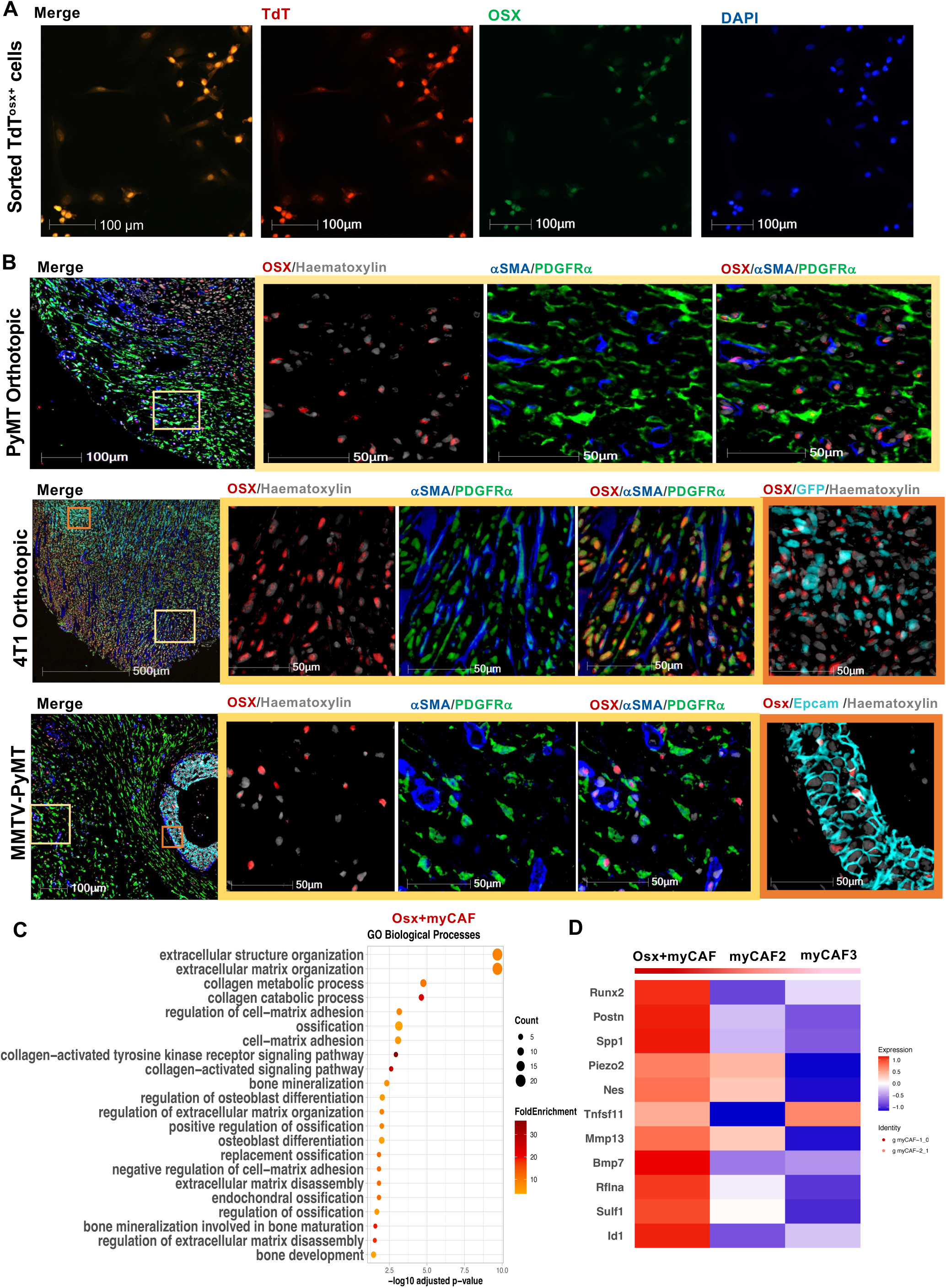
Tumor-infiltrating Osx^+^myCAF retain an osteolineage signature. **A,** Representative images of TdT^+^OsteoLin stromal cells sorted from PyMT-BO1-GFP^+^ tumors in OsxCre;TdT mice and plated onto coverslips for IF analysis. Osx staining (green), DAPI (blue) and TdT^+^ cells are shown in red. **B,** Representative images of multiplex Immunohistochemistry (mIHC) from orthothopic PyMT, orthothopic 4T1 and spontaneous MMTV-PyMT breast tumors (n=3/group) stained for Osx (red), Hematoxylin (gray), αSMA (blue), PDGFRα (green) and GFP (4T1, cyan) or EpCAM (MMTV-PyMT, cyan). **C,** Gene ontology (GO) Pathway Analysis from murine scRNA-seq data showing enrichment of osteogenic pathways in the Osx^+^myCAFs. Dot size indicates the number of genes in each pathway, while dot color represents the proportion of the top 50 differentially expressed genes (DEGs). **D,** Heatmap illustrating the expression of osteogenic markers among myCAFs populations.

To assess Osx expression in murine tumor tissues, we performed multiplex immunohistochemistry (IHC) in the orthotopic PyMT, triple-negative 4T1 BC models, and the spontaneous MMTV-PyMT model. For each antibody used, images were given an arbitrary color with the deconvolution algorithm provided by HALO software under the supervision of a trained pathologist. Osx nuclear staining was observed in subsets of PDGFRα⁺ and αSMA⁺ cells across all tumor models. Additionally, in the 4T1 BC and spontaneous MMTV-PyMT models, Osx staining was also detected in a few GFP⁺ and EPCAM⁺ cancer epithelial cells (Fig. 3B). However, in the orthotopic PyMT model, tumor cells were not fluorescently tagged and lacked EPCAM expression, precluding the assessment of Osx staining in this compartment. To further characterize the nature of Osx^+^myCAFs, we compared the gene signature of the three myCAF subclusters identified in the scRNAseq. GO analysis revealed enrichment in pathways related to bone mineralization, osteoblast differentiation, ossification, and ECM organization/remodeling in the Osx^+^myCAF subset (Fig. 3C), and high expression of osteogenic markers, such as *Runx2, Postn, Spp1 (Opn), Piezo2, Nes, Tnfsf11, Mmp13, Bmp7, Rflna, Sulf1, and Id1* (Fig. 3D). Altogether, these results indicate the existence of an Osx^+^myCAF subset retaining a strong osteogenic signature that is not found in myCAF2 and myCAF3 subsets.

### Osterix expression in CAFs drives tumor growth

To understand if Osx is merely a marker of an osteolineage-derived myCAF or directly contributes to their pro-tumorigenic function, we ectopically expressed Osx in murine mammary fibroblasts (MMFOsx^+^) or CAFs derived from PyMT-MMTV spontaneous tumors (FVB background) using pLenti-GFP-puro reporter vector (Suppl. Fig. 4A). MMFs or CAFs expressing empty vectors were used as controls (MMFctr and CAFctr). Infection efficiency was confirmed by detecting the GFP signal via flow cytometry and Osx transcripts by qPCR (Suppl. Fig. 4B-D). Next, we generated 3D spheroids using PyMT-mCherry tumor cells alone or together with MMFctr or MMFOsx^+^ (Fig. 4A). Tumor growth was quantified in real-time using live-cell fluorescence microscopy with a red channel for mCherry detection. Similarly to mice co-injected with TdT^+^OsteoLin cells, PyMT spheroids grew better in the presence of MMFOsx^+^ compared to MMFctr or PyMT alone (Fig. 4B-C). Notably, PyMT formed aggregates when cultured with MMFs, a phenomenon known to facilitate tumor progression (32), and the MMFOsx^+^ group displayed significantly more tumor aggregates compared to controls (Fig. 4D).

**Figure 4.**
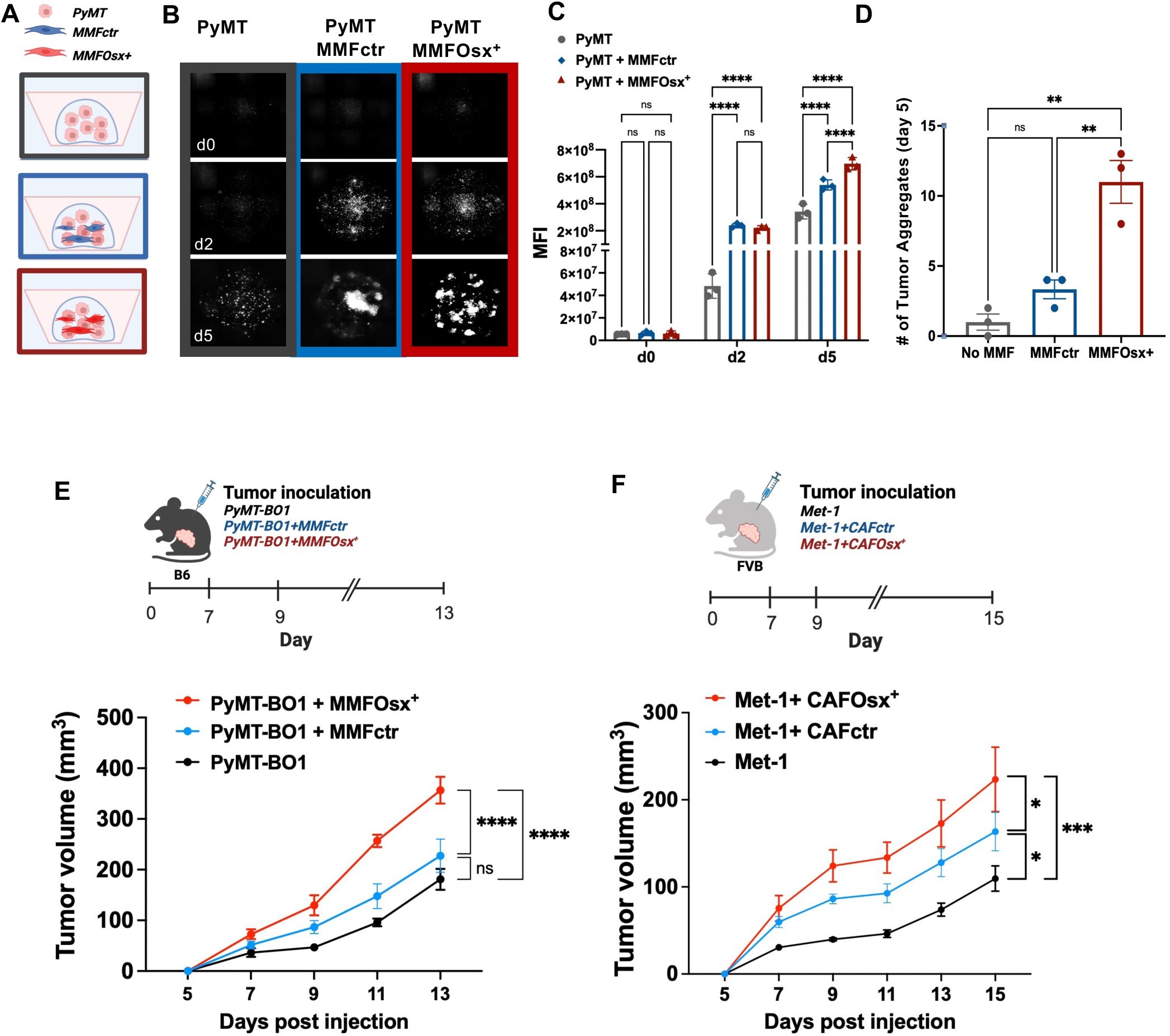
Osx expression in CAFs drives tumor growth. **A-D,** (A) Experimental scheme of PyMT-mCherry^+^ spheroids cultured alone or with MMFctr or MMFOsx^+^. (B) Representative images of spheroids taken with a fluorescent microscope on days 0, 2, and 5 after plating. n = 3/group. (C) Quantification of (B) as maximum fluorescence intensity (MFI) over time, and (D) number (#) of tumor aggregates formed at day 5 in each group. **E,** Schematic of PyMT-BO1-GFP^+^ tumor cell co-injections with MMFctr or MMFOsx^+^ at a 1:1 ratio into C57B/6 WT female mice (top) and tumor growth determined by caliper measurements of over time. Tumor cells alone were used as a control. (n = 6/group). **F,** Schematic of Met-1 tumor cell co-injections with CAFctr or CAFOsx^+^ at a 1:1 ratio (top) into FVB WT female mice and tumor growth determined by caliper measurements over time. Tumor cells alone were used as a control (n = 6/group). Experiments were conducted in at least three independent replicates. Results are shown as mean +/- SD, Two-way ANOVA followed by Turkeýs multiple-comparison test was used to determine significance in C, E, and F. One-way ANOVA followed Turkeýs multiple-comparison test was used to determine significance in D (* P≤0.05, **P≤0.01, ****P≤0.0001).

Next, we employed an *in vivo* model by co-injecting PyMT tumor cells either alone or with MMFOsx^+^ or MMFctr into eight-week-old WT female mice. MMFOsx^+^ significantly enhanced tumor growth compared to MMFctr (Fig. 4E). Similar results were obtained using the Met-1 BC cell line co-injected with Osx^+^CAFs into FVB female mice (Fig. 4F). These results demonstrate that Osx exerts pro-tumorigenic functions when expressed in CAFs, revealing its active role beyond being a marker of osteolineage-derived cells.

### MMP13 is required for Osx^+^myCAF pro-tumorigenic effects

In osteoblasts, Osx directly regulates the expression of MMP13, a matrix-degrading enzyme crucial for ECM remodeling and bone turnover (33). Given the similarities with osteoblasts, we investigated whether MMP13 contributes to Osx⁺myCAF pro-tumorigenic effects. First, we confirmed that MMP13 was highly expressed in both Osx⁺myCAFs and MMFOsx⁺ (Fig. 5A,B). Next, we cultured PyMT-BO1 tumor cells with conditioned media (CM) collected from either MMFctr or MMFOsx⁺. Crystal violet staining and MTT assay showed increased numbers of tumor cells when exposed to MMFOsx⁺ CM, forming larger and more elongated clusters compared to MMFctr CM or tumor cells alone (Fig. 5C-E). Notably, treatment with an MMP13 inhibitor (MMP13i CL-82198 20 µM), which specifically binds the S1 pocket of the enzyme and impairs its function, inhibited tumor cell growth and the morphological alterations induced by MMFOsx⁺ CM (Fig. 5F,G). In contrast, no differences in cell numbers or morphology were noted when the tumor cells were cultured in normal media, with or without MMP13i (Fig. 5F,G). Similarly, mCherry-labeled PyMT cells cultured with MMFOsx⁺ in a 3D spheroid model showed a marked reduction in tumor cluster size when treated with the MMP13i (20 µM) (Fig. 5H,I). We did not observe any direct effects of MMP13i on the tumor cells or the MMFs when cultured alone (Fig. 5J).

**Figure 5.**
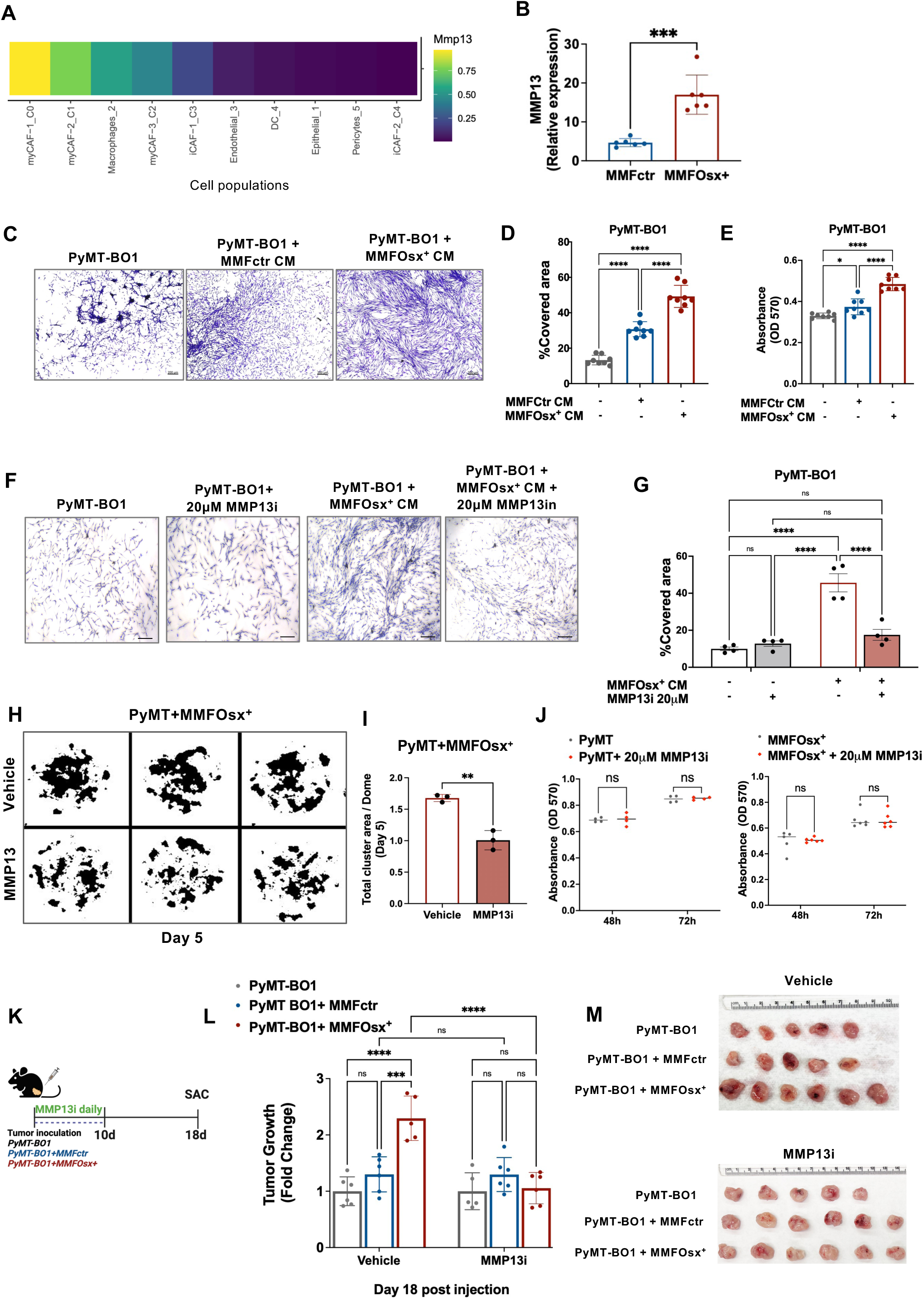
MMP13 is required for Osx^+^myCAF pro-tumorigenic effects. **A,** Heatmap illustrating MMP13 expression based on scRNA-seq data across indicated populations in orthotopic PyMT-BO1 tumors from Osx-Cre;TdT mice. **B,** qPCR analysis showing *Mmp13* transcripts in MMFOsx^+^ and MMFctr. **C-E,** Representative images of crystal violet staining of PyMT-BO1-GFP^+^ tumor cells after 48 hours in 1% FBS control media, or conditioned media (CM) collected from MMFctr and MMFOsx^+^ (C, Bar=200μM). (D) Quantification of area occupied by tumor cells from (C). (E) Number of viable tumor cells in control media, CM from MMFctr or MMFOsx^+^ determined by MTT colorimetric assay. **F-G,** (F) Representative images of crystal violet staining of PyMT-BO1-GFP^+^ tumor cells treated for 72 hours with control media or CM from MMFOsx^+^, in the presence or absence of MMP13i (20 μM, Bar=200μM). (G) Quantification of area occupied by the tumor cells in (F). **H-I,** Representative z-stack images of PyMT spheroids cultured with MMFOsx^+^ and treated with vehicle or 20 µM MMP13i for 5 days (H). Images were converted to 8-bit black-and-white format, and cluster areas were analyzed using ImageJ with a threshold range of 0–73. (I) Quantification of cluster area was performed based on thresholded images in (H). **J,** Number of viable tumor cells or MMFOsx^+^ treated with vehicle or 20 µM MMP13i for 48 and 72hrs determined by MTT colorimetric assay. **K-M,** (K) Schematic of MMP13i (0.5mg/mouse) or vehicle treatment of PyMT-BO1-GFP^+^ tumor cell co-injections with MMFctr or MMFOsx^+^ into the mammary fat pad of WT female mice. (L) Tumor volume measured by caliper with data represented as fold change from day 7 and (M) harvested tumors. Results are shown as mean +/- SD. Unpaired Student T-test was performed to determine significance in B and I. One-way ANOVA followed by Turkeýs multiple-comparison test was used to assess significance in D and E. Two-way ANOVA followed by Turkeýs multiple-comparison test was used to determine significance in G, J, and L. (* P≤0.05, **P≤0.01,***P≤0.001, ****P≤0.0001).

To assess the effects of MMP13i *in vivo*, PyMT-BO1 tumor cells were injected alone or with MMFOsx⁺ or MMFctr into the MFP of eight-week-old WT female mice. Mice were treated with MMP13i (0.5 mg/mouse) or vehicle for 10 consecutive days starting on the day of tumor injection and sacrificed 18 days post-tumor inoculation (Fig. 5K). While no anti-tumor effects were observed in mice co-injected with MMFctr receiving MMP13i, a significant reduction in tumor burden was detected in the MMFOsx⁺ group upon MMP13i treatment (Fig. 5L,M). These findings indicate that osteoprogenitor-derived Osx⁺myCAFs promote tumor progression, at least in part, through the release of MMP13.

### myCAFs with an osteolineage signature are detected in human breast cancer tissues

To investigate whether Osx^+^myCAFs are detected in human BC tissues, we analyzed CAF clusters from scRNA-seq data generated by two independent studies to obtain an integrated data set comprising 40 samples of TNBC, ER+, and HER2+ BC cases (Fig. 6A; (18, 34)). UMAP dimensionality reduction and graph-based clustering yielded the 6 previously reported BC CAF subsets: Vasculature CAFs (vCAFs) marked by *MCAM* and *NOTCH3*, myCAFs marked by *POSTN* and *TAGLN*, iCAFs marked by *CXCL12*, and *COL14A1*, antigen-presenting CAFs (apCAFs1 and apCAFs2) marked by *CD74* and MHCII genes such as *HLA-DRA*, and dividing CAFs (dCAFs), marked by *MKI67* and *TOP2A* (Fig.6B, Suppl. Fig.5A). Based on *PDGFRα* and *PDGFRβ* expression in the murine Osx^+^myCAF subset, we selected clusters positive for both markers (Fig. 6C,D). Re-clustering revealed one h-iCAF population and three transcriptionally distinct myCAF subpopulations referred to as h-myCAF#1–3 (Fig. 6E,F, Suppl. Fig.5B). To identify the human ortholog of mouse Osx^+^myCAF, we mapped the top 50 differentially expressed genes (DEGs) and 10 curated osteolineage markers from murine Osx^+^myCAFs onto the human dataset using Gene Set Enrichment Analysis (GSEA). We found an equivalent population in the h-myCAF1 cluster showing elevated expression of osteogenic markers *RUNX2, POSTN, SULF1, MMP13, SPP1, and TGFB1* (Fig. 6G). This osteolineage signature was either not positively enriched or showed negative enrichment in the other h-myCAF2 and h-iCAF populations (Fig. 6F-H, Suppl. Fig. 6A). Of note, the murine Osx^+^myCAF signature partially overlapped with the h-myCAF3 cluster (Suppl. Fig. 6B). However, h-myCAF3 showed enrichment in immunogenic genes, such as *RSAD2, ISG15, IFIT1*, and *IFIH1,* and pathways associated with interferon response and production (Fig. 6F, Suppl. Fig. 6C), which were not detected in the murine Osx^+^myCAFs (Suppl. Fig. 6D).

**Figure 6.**
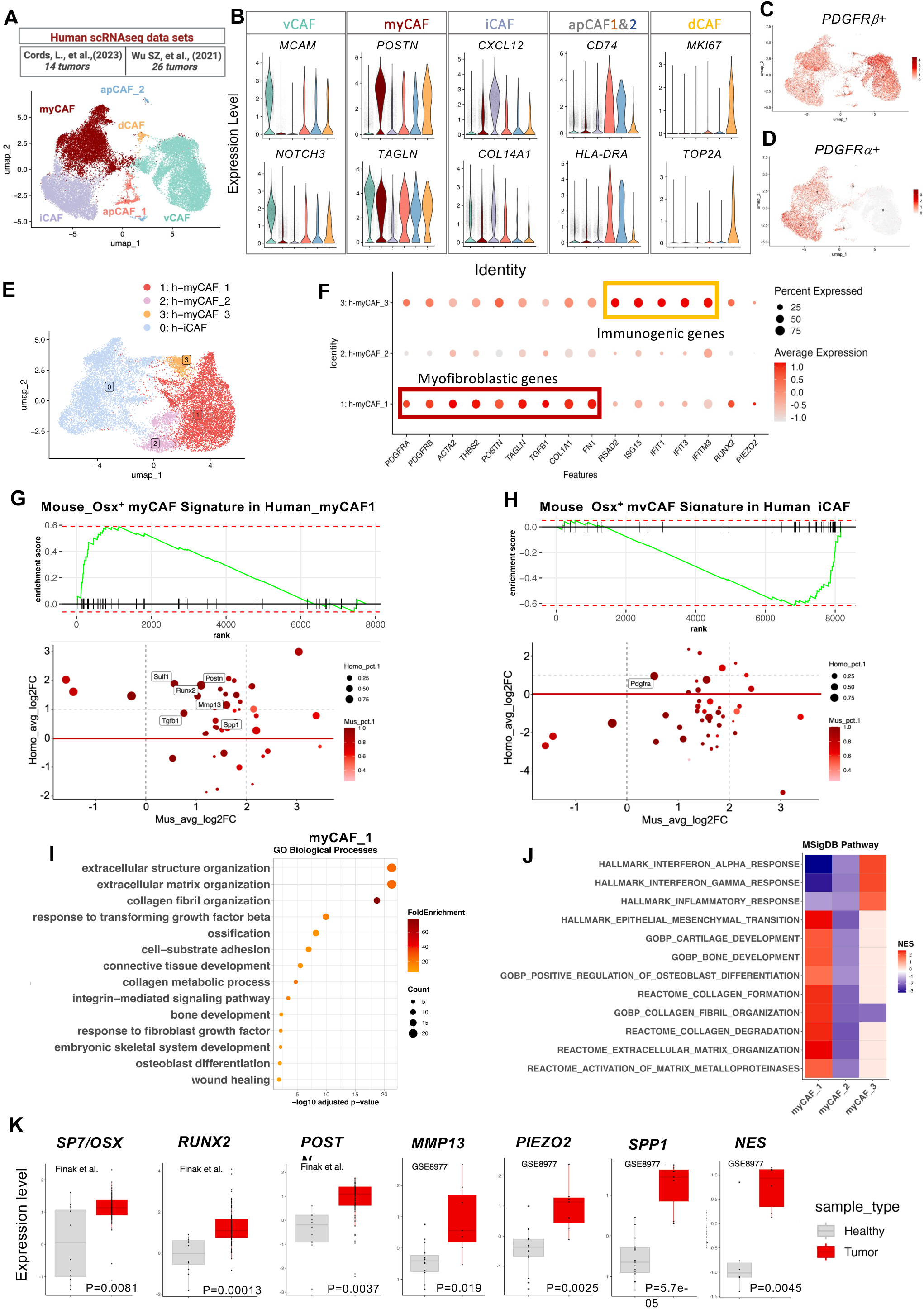
Human myCAF1s express an osteolineage signature. **A,** Schematic of two human breast cancer scRNA-seq data sets (top) used to identify CAF clusters by UMAP. Data shows 6 clusters (C0: vCAF, C1: myCAF, C2:iCAF, C3:apCAF_1, C4: apCAF_2, C5: dCAF). **B,** Violin plots of human breast cancer CAF subsets showing 2 markers for each subset. **C-E,** Featured plots of CAF subsets expressing *PDGFRβ* (C) and *PDGFRα* (D) genes. (E) UMAP of *PDGFRβ / PDGFRα* h-CAF subclusters (C0: h-iCAF, C1: h-myCAF1, C2: h-myCAF2, C3: h-myCAF3). **F,** Dot plots showing expression of CAF marker genes across h-myCAF1-3, with dot size indicating the percentage of cells expressing indicated gene in each cluster and dot color representing expression levels. **G-H**, GSEA (top), and scatterplot (bottom) showing overlap of murine Osx^+^myCAF signature genes with the h-myCAF1 (G) and h-iCAF (H) clusters. **I-J,** (I) Gene Ontology (GO) pathway analyses showing enrichment of genes involved in ossification, bone development, ECM remodeling, and collagen organization in h-myCAF1. (J) Heatmap based on GSEA pathway analysis showing ECM-related signaling pathway across h-myCAF clusters. **K**, Human microarray dataset analyses (Finak et al. or GSE8977) showing expression levels of osteoblast related markers in healthy versus tumor stroma from breast cancer tissues. Unpaired Student T-test was performed to determine the significance in K.

Similarly to the Osx^+^myCAFs, GSEA analysis and GO enrichment of the top 50 DEGs, further demonstrated that the h-myCAF1s were enriched in genes associated with ECM remodeling, collagen reorganization, and metalloproteinase activation (Fig. 6I). Notably, this population was also enriched in bone-related pathways, including ossification, bone development, cartilage development, and osteoblast differentiation (Fig. 6I,J). Expression of osteoblastic genes in the human tumor stroma was also detected in two microarray datasets (Finak Dataset (35) and GSE8977 (36)) comprising tissues from both healthy and malignant breast samples. SP7*, RUNX2, POSTN, SPP1, PIEZO2, NES, and MMP13* were increased in the human BC stroma compared to the healthy counterpart (Fig. 6K).

Collectively, these findings demonstrate the presence of the h-myCAF1 subset in BC tissues, with strong overlap with osteolineage-derived Osx^+^myCAF in mice.

### Osterix is expressed in human breast cancer stroma and is associated with poor prognosis

To assess Osx expression in hCAFs, we performed mIHC on tissue microarrays containing triple-negative BC (TNBC, N=10) and estrogen receptor-positive (ER⁺, N=8) BC samples. Pan-cytokeratin (panCK) was used to identify epithelial/tumor cells, while myCAFs were identified by αSMA and PDGFRα staining. Like the murine BC models, we detected a strong nuclear Osx staining in subsets of αSMA^+^/PDGFRα^+^ populations in both ER+ and TNBC tissues (Fig.7A).

**Figure 7.**
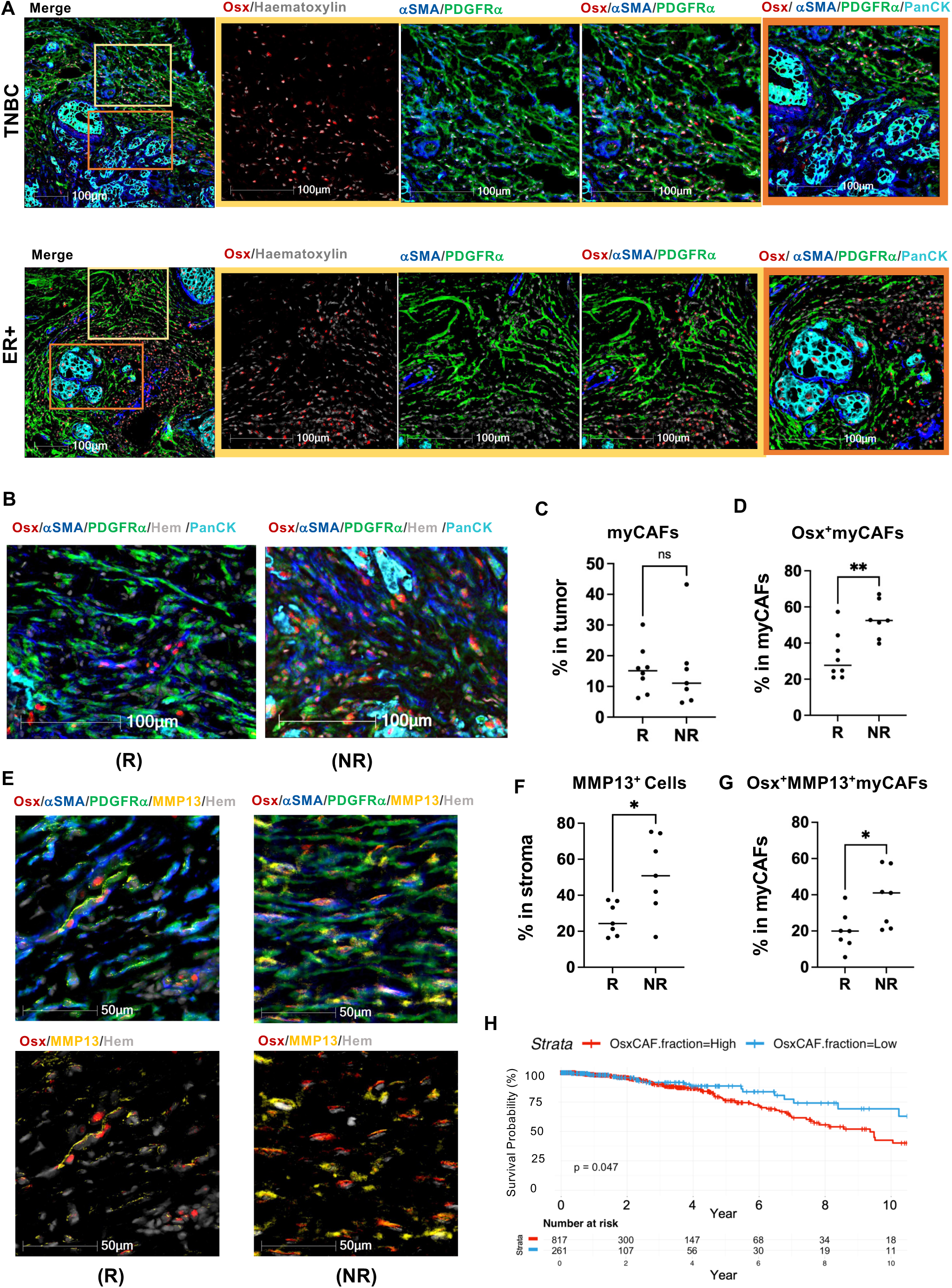
Osx^+^myCAFs are present in human breast tumors and correlate with poor prognosis. **A,** Representative mIHC images of triple-negative (n=10), and ER^+^ (n=8) human breast cancer tissues stained for Osx (red), Hematoxylin (gray), αSMA (blue), PDGFRα (green) and panCK (cyan). The yellow inset highlights the stromal area, and the orange inset highlights the tumor area. **B,** Representative mIHC images of TNBC patient biopsies at diagnosis before receiving neoadjuvant standard of care. Based on annotated therapeutic response, patients were classified as responders (R=8) or no responders (NR=7). Tissues were stained for Osx (red), Hematoxylin (gray), αSMA (blue), PDGFRα (green), and panCK (cyan). **C,** Quantification of myCAFs (PDGFRα^+^/αSMA^+^ cells) within the stroma (PanCK^neg^ cells) in responders vs. non-responders. **D,** Quantification of Osx^+^myCAFs (Osx^+^ PDGFRα^+^ αSMA^+^ cells) within the stroma (PanCK^neg^ cells) in responders vs. non-responders. **E,** Representative mIHC performed as described in (B) on the same tissue samples, with the addition of MMP13 antibody. **F,** Quantification of MMP13^+^ cells within the stroma (PanCK^neg^ cells) in responders vs. non-responders. **G,** Quantification of MMP13^+^Osx^+^myCAFs (PDGFRα^+^/αSMA^+^ cells) out of total myCAFs in responders vs. non-responders. **H,** Kaplan-Meier plots of overall survival for breast cancer patients segmented by high (red line) or low (blue line) abundance of orthologous Osx^+^myCAF using the TCGA-BRAC database. Results are shown as mean +/- SD. Unpaired Student T-test was performed to determine the significance in C,D, F, G, and H. (* P≤0.05, **P≤0.01).

We then analyzed 15 clinically annotated BC biopsies from TNBC patients at the time of diagnosis, before receiving standard-of-care neoadjuvant therapy (chemotherapy plus anti-PD1). After completion of neoadjuvant therapy, the patients were classified based on the Residual Cancer Burden (RCB) index. Seven patients were classified as RBC II and III, having moderate to large amounts of remaining tumor cells (non-responders; NR), while eight as RCB 0 because achieved a pathological complete response (responders; R). Tissues were subjected to mIHC for expression of Osx, MMP13, αSMA^+^/PDGFRα^+^ myCAF, panCK epithelial marker, and hematoxylin for nuclear staining (Fig 7B,E; representative images) followed by quantification using the Halo software. Strikingly, the proportion of h-myCAFs positive for Osx was significantly higher in the non-responders, while the overall proportion of h-myCAFs (αSMA⁺/PDGFRα⁺) was similar between responders and non-responders (Fig. 7B-D). The increase in h-Osx⁺myCAFs was also accompanied by elevated MMP13 expression (Fig.7E-G), suggesting a potential link between Osx-driven ECM remodeling and therapy resistance.

Finally, we analyzed the bulk RNA-seq data from TCGA-BRCA dataset (37) to evaluate whether the abundance of h-Osx^+^myCAF could predict patient outcomes. To infer the proportion of h-Osx^+^myCAF, we applied CIBERSORTx (38) to separate mixed cell populations inside the bulk RNA-seq data. Patients were then classified as h-Osx^+^myCAF-high (above the lower quantile) or h-Osx^+^myCAF-low (equal to or below the lower quantile). Our analysis revealed that higher h-Osx^+^myCAF proportions were significantly associated with reduced 10-year survival in BC patients (p = 0.047; Fig. 7H).

## Discussion

In this study, we identified a specialized subset of myCAFs originating from BM-derived Osx^+^ osteoblast progenitors, involved in BC progression through ECM remodeling and production of MMP13. Notably, the presence of Osx^+^myCAFs is associated with poor therapeutic response and reduced overall survival of breast cancer patients. These findings provide new insights into the origin of a specialized myCAF subset and suggest a potential pathway to target tumors enriched with h-Osx^+^myCAFs.

Despite the identification of multiple specialized CAF populations, it is not clear whether CAF subsets originate from distinct progenitors or derive from a common precursor that adapts to local cues to acquire specific phenotypic traits and functions. We find that BM resident Osx^+^ osteolineage cells, known to differentiate into bone-forming osteoblasts, can be mobilized to extraskeletal tumors and give rise to a unique Osx^+^myCAF subset in the TME, with increased pro-tumorigenic effects compared to Osx^neg^CAF populations. Previous studies have reported circulating CD15⁻CD34⁻Osteocalcin⁺ cells (Osteocalcin is a marker of osteoblasts and Osx downstream target) linked to BC metastases (39). Similarly, we detect osteolineage-derived cells in circulation in tumor-bearing mice, suggesting analysis of the osteolineage marker Osx could be useful for diagnostic purposes. Further investigation is still needed to determine whether circulating Osx⁺myCAFs are present in patients and to assess their clinical relevance across disease stages.

Interestingly, unlike other previously reported BM-derived CAFs (20), Osx⁺myCAFs express PDGFRα and have ECM remodeling properties, rather than pro-inflammatory functions. Like bone-residing Osx^+^ osteoblasts, Osx⁺myCAFs express key bone-associated markers, including *Runx2*, *Msx1*, osteopontin (*Spp1*), periostin (*Postn*), and *Mmp13* (33, 40). Expression of osteogenic genes and ECM remodeling properties are also detected in a specialized subset of human myCAFs. Notably, we have identified a 60-gene osteolineage signature comprising common, highly expressed bone-related genes in murine and human Osx^+^myCAFs correlating with poor BC survival, indicating that these cells retain their origin identity. Interestingly, Osx^+^ osteolineage cells directly isolated from the bone fail to directly support BC growth, suggesting that this feature is acquired in the TME and modulated by tumor-derived signals. These results demonstrate that bone-resident Osx^+^ osteolineage cells can acquire fibroblastic features at different tumor sites. It also cannot be excluded that bone-derived Osx^+^ osteolineage cells contribute to fibroblast diversity in other tissues or diseases. Expression of *Runx2*, a key osteogenic gene expressed in the Osx^+^myCAF subset, has been shown to drive the generation of pathological alveolar fibroblasts and ECM deposition in pulmonary fibrosis (41). However, whether these fibroblasts derive from an Osx^+^ bone population was not assessed.

For decades, Osx expression was solely thought to be confined to the bone. With the use of OsxCre;TdT lineage tracing mouse model, the presence of TdT^+^OsteoLin cells was detected outside of the skeleton, and during cancer progression within the TME (29). Surprisingly, most of these cells constituted tumor-infiltrating CD45^+^ immune populations derived from a rare Osx^+^ hematopoietic stem cell that loses Osx expression upon maturation (29). In another study, constitutive activation of NF-κB inducing kinase (NIK) under the OsxCre promoter drove spontaneous extra-skeletal soft tissue sarcomas (42). The staining profile and Cre expression suggested a mesenchymal origin, though it was unclear whether the tumor arose from BM-derived Osx⁺ cells mobilized outside the bone. Osx has also been detected in BC cells. Over-expression and knockdown studies in BC cell lines demonstrated Osx involvement in metastatic dissemination (43). We now show that Osx is also expressed in a specialized subset of myCAFs, representing about 1% of cells in the tumor mass and 20% of PDGFRβ^+^ stromal populations. By combining the OsxCre;TdT lineage tracing approach with scRNAseq analyses, we identified 3 myCAF subclusters, of which only myCAF1 is highly enriched in TdT^+^ cells and expresses osteogenic markers. We report similar findings in the scRNAseq human data sets, where out of 3 h-myCAF subclusters only h-myCAF1 shows an osteolineage signature. Our findings are in line with the recent identification of highly specialized myCAFs, including contractile myCAFs enhancing ECM stiffness, and ECM-remodeling myCAFs, secreting collagens and MMPs to reshape the TME (44). Given the similarities in functionality and gene expression profile with our Osx^+^myCAFs, it is likely that contractile and ECM-remodeling CAFs might derive from a BM Osx^+^ osteoprogenitor. While we cannot detect Sp7/Osx in CAF clusters by scRNAseq, possibly due to limited sequencing depth or rapid Osx/Sp7 mRNA degradation, we confirmed Osx nuclear localization by mIHC in subsets of PDGFRα⁺ and αSMA stromal populations in murine and human BC tissues. Notably, Osx⁺myCAFs are abundant at diagnosis in TNBC patients resistant to standard-of-care, yet the overall percentage of myCAFs remains similar between responders and non-responders. While myCAFs are linked to poor prognosis, our findings suggest that analyses of myCAF subclusters at the time of diagnosis may be used to determine the therapeutic response.

Osx is not just a marker of osteoprogenitor-derived myCAFs but also drives their tumor-promoting function. While its role in osteoblast differentiation is well established and its expression in tumor cells promotes invasion (27, 43), no prior work has described Osx function in CAFs. Osx regulates expression of ECM remodeling proteins, including collagens and MMPs. MMP13, a direct Osx target gene, is highly expressed in Osx^+^myCAFs compared to the other myCAF subclusters. In BC, MMP13 is linked to aggressiveness and poor prognosis (45, 46) by modulating collagen degradation, and the release of ECM-stored bioactive molecules to support EMT, angiogenesis, and immune modulation. We find that conditioned medium from MMFOsx⁺ induces pronounced morphological and functional changes in BC cells, such as acquisition of an elongated and more spread morphology, and ability to grow in clusters. Similar findings are observed in our 3D cultures of tumor cells with MMFOsx^+^ showing a significant increase in tumor aggregates compared to MMFctr or tumor cells alone. These results are in line with the described changes in BC cell phenotype following exposure to CAF-CM (47). Importantly, MMP13i completely rescues the morphological changes in the tumor cells driven by MMFOsx^+^, most likely through inhibition of ECM remodeling and preventing exposure of bioactive signals. Furthermore, MMP13i significantly reduces tumor growth in mice co-injected with MMFOsx⁺ but not MMFctr. These findings align with evidence that MMP13 knockdown in myoepithelial cells reduces tumor invasion in 3D models (48) and that stromal MMP13 drives DCIS-to-IDC transition (49). Expression of MMP13 has also been detected in a subset of BM-derived myofibroblasts in skin carcinoma promoting tumor invasion (50). We find that MMP13 is primarily expressed in Osx^+^myCAFs from murine and human BC tissues. Importantly, MMP13^+^Osx^+^myCAFs are increased in biopsies from TNBC patients not achieving a pathological complete response to a neoadjuvant standard-of-care. These findings have important clinical implications, as they suggest that MMP13 is a critical regulator of Osx^+^myCAF pro-tumorigenic properties and its targeting could be beneficial to patients with elevated Osx stromal expression. However, given that Osx can also control the expression of several collagen genes and other ECM-related proteins, a multi-targeted approach might be needed to achieve a highly efficient anti-tumor response in the clinic.

In conclusion, we have demonstrated the existence of a subset of Osx^+^myCAFs, originating from a bone Osx+ osteoblast progenitor, with osteolineage features that distinguish its phenotype from other myCAF subsets, and whose presence correlates with reduced therapeutic response and survival in BC.

## Materials and Methods

### Animals

Because the breast cancer cell lines used in this study were obtained from female mice, we have restricted our analyses to females. Wild-type (WT) C57BL/6, FVB/N, BALB/c, B6.SJL-Ptprca Pepcb/BoyJ (CD45.1 B6, JAX #002014), Sp7Cre (B6.Cg-Tg(Sp7-tTAtetO-EGFP/Cre), and TdT (B6.Cg-Gt(ROSA)26SorTm9(CAG-tdTomato)Hze/J were purchased from The Jackson Laboratory at 6-8 weeks of age. After delivery, mice were allowed to acclimatize to the new environment for at least 2 weeks. Animals were housed in a pathogen-free animal facility at Washington University (St. Louis, MO) with a 12-h light/12-h dark cycle and 20∼23 °C and 40∼60% humidity housing conditions. Sp7Cre mice, which harbor a tetracycline-responsive Osx promoter driving Cre (Osx-cre), were crossed with TdT reporter mice to generate Osx-cre;TdT mice. To suppress Cre expression, 200ppm doxycycline (doxy) was added to the chow (Test Diet #1816332–203, Purina, MO, USA) and administered to specific groups of mice until weaning (P25); thereafter, pups were transitioned to standard rodent chow. All experiments were performed according to protocols approved by the Institutional Animal Care and Use Committee at Washington University (Protocol ID: 2022-0315).

### Cell lines

Polyoma middle tumor-antigen murine mammary tumor cells (PyMT, C57BL/6), mCherry-conjugated PyMT (from D. DeNardo), bone-trophic GFP- and firefly-luciferase-conjugated PyMT-BO1 (from Dr K. Weilbaecher), H2B-mApple-Thy1.1- and fluc-conjugated PyMT-BO1 (from Dr S.A. Stewart), GFP- and fluc-conjugated 4T1 murine mammary tumor cells (BALB/c), Met-1 breast cancer cell line (FVB), mammary fibroblasts (MMF, C57BL/6) and immortalized CAFs (FVB) were cultured at 37°C with 5% CO2 in complete media DMEM supplemented with 10% heat-inactivated FBS, 100 μg/ml streptomycin, 100 IU/ml penicillin, and 1 mM sodium pyruvate. All cell lines were tested for *Mycoplasma* every 2 months. Aliquots for each cell line were used for a maximum of one month after the initial thaw.

### Tumor Cell Inoculation

Tumor cells (10^5^ cells) resuspended in 1:1 PBS/Matrigel ratio (Corning 354234) at a final volume of 50 µl, were injected into the mammary fat pad of 6-8 weeks female mice. For co-injections, 10^5^ PyMT or PyMT-BO1 cells were inoculated with sorted CD45^neg^ TdT^+^OsteoLin cells, MMFctr, or MMFOsx^+^ at a 1:1 or 1:2 stroma-tumor ratio. Similarly, 10^5^ Met-1 tumor cells were co-injected in a 1:1 ratio with control or Osx^+^CAFs in FVB female mice. When indicated, the MMP13 inhibitor CL-82198 (MedChem Express) dissolved in 10% DMSO and diluted in 40%PEG300 and 50% PBS was injected intravenously (0.5 mg/mouse) to tumor-bearing mice for 10 consecutive days. Tumor measurements began 7 days post-injection and were conducted every other day using a caliper. Tumor volumes were calculated using the formula: V = 0.5 (length [mm] x width [mm]^2^).

### Bone Marrow Transplantation

Recipient C57BL/6 CD45.1 mice received two doses of 400 cGy 4 hours apart, using an X-ray irradiator (XRAD 320), followed by transplantation of BM cells collected from at least two CD45.2 OsxCre;TdT donor mice. Briefly, CD45.2 OsxCre;TdT donor mice were sacrificed by CO₂ inhalation, and femurs, tibias, and ilia were harvested under sterile conditions. Bones were placed in 0.5ml conical tubes perforated at the bottom and inserted into larger 1.5ml tubes. BM was flushed using pulsed centrifugation, transferred into clean 1.5ml tubes, and resuspended in cold, sterile, serum-free 1X HBSS. 200 μL containing 5×10⁶ BM cells were administered via retro-orbital injection into each CD45.1 recipient mouse. Mice were given sucrose water for a week and monitored for the following 2 weeks for signs of radiation sickness or weight loss. Six weeks after irradiation, mice were used for indicated experiments.

### Multiparametric Flow Cytometry

Tumors were digested in serum-free media with 2 mg/ml collagenase type I and 2 U/ml DNaseI for 30 min at 37°C. The cell suspension was filtered through 70-μm nylon strainers and washed twice in PBS with 2% FBS. Red blood cells were removed using lysis buffer (Sigma-Aldrich), followed by an additional wash. Cells were then stained in PBS with 0.5% FBS using anti-mouse antibodies. For blood analyses, blood was collected from mice via cardiac puncture. Red blood cells were removed by two 10-minute incubations with lysis buffer, followed by an additional wash to ensure complete lysis. The remaining cells were then processed for flow cytometric staining and subsequently acquired for analysis. Acquisition was performed on a BD LSRFortessa X-20 Cell Analyzer with Diva software (BD), and data were analyzed using FlowJo 10.9.0 (Tree Star).

### Immunofluorescence staining and confocal imaging of orthotopic tumors

Tumors were fixed in 4% paraformaldehyde (PFA) at 4°C overnight. Fixed specimens were then dehydrated in 30% sucrose solution and cut into 50 μm-thick sections at the cryostat (Leica). Staining on free-floating sections was performed. Cryosections were blocked for 4 hours in 5% BSA solution with 0.5% Triton X-100 and stained with primary antibodies for 48h at 4°C. Secondary staining was performed at room temperature for 2 h. Sections were then mounted on Superfrost glass slides (Fisher Scientific) and embedded in Prolong Glass anti-fade mounting media (Thermofisher). Sections were covered with 1.5H high precision cover glass (Marienfeld Superior) and let dry overnight at room temperature before imaging. Confocal imaging of tumor cryosections was carried out using a Zeiss LSM880 airyscan inverted confocal microscope, with a 40X/NA1.4 oil immersion objective. Images were acquired at 2048X2048-pixel resolution, 10/30μm-thick *z*-stack z-step=1μm, line averaging=2, using ZEN Black (ZEISS Efficient Navigation) software (Zeiss). Maximal projections were rendered in ImageJ (version 2.3.0).

### Histology and Immunohistochemistry

Freshly isolated mouse primary tumors were fixed in 10% neutral-buffered formalin (DiRuscio & Associates, Inc.) for 24 h. Tissues were paraffin-embedded and sectioned 5μm thick by the histology core of the Washington University Musculoskeletal Research Center.

Human tissue microarrays were prepared by The St. Louis Breast Tissue Registry (funded by The Department of Surgery at Washington University). Data and tissues were obtained following the guidelines established by the Washington University Institutional Review Board (IRB #201102394) and WAIVER of elements of Consent per 45 CFR 46.116 (d). All patient information was deidentified before sharing it with investigators. Clinically annotated diagnostic biopsies from TNBC patients were obtained from Department of Pathology at Washington University. Patients were subjected to neoadjuvant standard-of-care regimen (Keynote522) consisting of pembrolizumab (200 mg) every 3 weeks plus paclitaxel and carboplatin weekly for the first 12 weeks, followed by evaluation of Residual Cancer Burden (RCB) score. Responders were selected based on RCB=0 and Non-Responders on RCB=II-III. Tissues from 8 responders and 7 non-responders were used for mIHC.

Tissues were automatically stained via the Bond Rxm (Leica Biosystems) following dewaxing and appropriate epitope retrieval. Immunostaining was chromogenically visualized using the Bond Polymer Refine Detection (#DS9800, Leica Biosystems) or the Bond Polymer Refine Red Detection (#DS9390, Leica Biosystems). Slides were mounted using Xylene-based Cytoseal (Thermo Fisher) or Vectamount (Vector Labs) as appropriate, scanned via a Zeiss AxioScan 7 microscope and IHC analyses performed using the HALO image analysis platform (Indica Labs, Deconvolution v1.1.1, Multiplex IHC v.3.2.3 algorithms).

### Lentivirus preparation and cell infection

The puromycin-resistant pLenti-puro ORF clone of Sp7 (mGFP-tagged – Origene MR226379L4) was used to ectopically express *Sp7* (*Osx*) in MMFs or immortalized CAFs. pLenti-puro (Addgene plasmid #39481) was used as a vehicle control. Briefly, HEK293T cells were transiently transfected with lentiviral accessory plasmids (VSV.G and psPAX2) and the designated transfer plasmid using TransIT-293 transfection reagent according to the manufacturer instructions (MIR2700, Mirus Bio LLC). The virus-containing cell supernatant was collected 48 hours after transfection, filtered through 0.45-μm filter, supplemented with 8 mg/mL protamine sulfate, and immediately added to target cells. Selection of cells expressing vehicle or Osx was performed in Dulbecco’s modified Eagle’s (DMEM) medium containing 2μg/ml puromycin and supplemented with 10% heat-inactivated fetal bovine serum, 100 IU/ml penicillin plus 100μg/ml streptomycin, and 1mM sodium pyruvate.

### MTT Assay of tumor cells exposed to CAF-conditioned media

MMFctr or MMFOsx⁺ cells were cultured in Dulbecco’s Modified Eagle Medium (DMEM) supplemented with 10% FBS, 1% sodium pyruvate, and 1% PenStrep. Once cells reached approximately 90% confluence, they were washed with PBS and incubated in low serum medium (DMEM containing 1% FBS, 1% sodium pyruvate, and 1% PenStrep). Conditioned media (CM) was collected after 24 hours. Tumor cells were seeded in 96-well plates (5 × 10³ cells/180μl per well) and treated with 100% CM from MMFctr or MMFOsx^+^. Tumor cells cultured in 1% FBS media were used as a control. After 48 hours, cell viability was evaluated using the MTT assay (Invitrogen) following the manufacturer’s protocol. In some experiments, tumor cells were cultured for 72 hours with or without MMFOsx⁺ CM, in the presence or absence of MMP13i (20 µM). MTT assay was also performed on tumor cells or MMFOsx^+^ alone cultured in DMEM supplemented with 10% FBS, 1% sodium pyruvate, and 1% PenStrep in the presence or absence of MMP13i.

### Crystal Violet Assay

Tumor cells cultured for 48-72 hours at 37°C in the presence of MMFctr or MMFOsx^+^ CM and treated with vehicle or 20 µM MMP13i (details described in MTT assay), were washed once with 1X PBS and fixed with cold methanol for 10 minutes at room temperature. The fixed cells were washed twice with 1X PBS to remove residual methanol and subsequently stained with a crystal violet solution for 10 minutes. Excess stain was removed by thoroughly rinsing the wells with distilled water. After air drying, stained cells were visualized under a microscope, and the covered area was calculated using ImageJ (version 2.3.0).

### 3D Spheroids culture

Three-dimensional spheroid cultures were established using Cultrex Organoid Qualified BME, Type 2 (R&D Systems, Inc., 3532-005-02). A total of 2.5 × 10⁴ mCherry⁺ PyMT cells, either alone or combined with MMFctr or MMFOsx⁺ at a 1:1 ratio, were resuspended in 20 µl of Matrigel. Domes were plated into a 6-well plate (three domes per well, per condition). The plate was placed in a tissue culture incubator at 37 °C with 5% CO₂ to allow the organoids to polymerize for 20 minutes. After polymerization, 2 mL of medium (DMEM supplemented with 10% FBS, 1% sodium pyruvate, and 1% PenStrep) was added to each well. In some experiments, 20 µM MMP13i was added to the media every two days with half media change, and the experiment was stopped on day 5. At least *n* = 3/group was used for statistical power analysis. mCherry^+^PyMT tumor cell growth was monitored by fluorescence imaging using a Nikon Eclipse Ti2 (temperature 37°C, 5% CO_2_, and a humid environment) and a 4x objective stitch to image the entire dome. Images were captured every 2 days using an ORCA-Flash4.0 LT3 Digital CMOS camera and processed via NIS-Elements (version 5.42.04) 64-bit software. Z-stacks were obtained to reconstruct the entire dome. Images were analyzed using NIS-Elements software to quantify fluorescence intensity and assess the number of clusters, while ImageJ (version 2.3.0) was used to measure the surface area of cell clusters.

### Real-time PCR analysis

Total RNA was extracted with TRIzol (Invitrogen, CA USA) and quantified on a ND-1000 spectrophotometer (NanoDrop Technologies). The cDNA was synthesized with 1 μg RNA using High-Capacity cDNA Reverse Transcription Kit (#4368814, Applied Biosystems, CA USA). The amount of each gene was determined using Power SYBR Green mix on 7300 Real-Time PCR System (Applied Biosystems). Cyclophilin mRNA was used as housekeeping control. Specific primers for mice were as follows: *Cyclophilin*, 5’-AGC ATA CAG GTC CTG GCA TC-3’ and 5’-TTC ACC TTC CCA AAG ACC AC-3’; *Osterix*, 5’-AAG GGT GGG TAG TCA TTT GCA-3’ and 5’-CCC TTC TCA AGC ACC AAT GG-3’; *MMP13* 5’-ACA GGG GCT AAG GCA GAA A-3’ and 5’-CGC TAA GGA AAG CAG AGA GG-3’,’; The relative quantification in gene expression was determined using the 2^-ΔΔCt^ method.

### Single-cell RNA sequencing

GFP^neg^CD45^neg^Ter119^neg^CD31^neg^TdT^+^ or TdT^neg^ fractions were sorted from PyMT-BO1-GFP^+^ orthotopic tumors eleven days post-inoculation in OsxCre;TdT mice. Cells were resuspended in PBS with 0.04% BSA at a final concentration of approximately 1000 cells/μl. cDNA was prepared after the GEM generation and barcoding, followed by the GEM-RT reaction and bead cleanup. Purified cDNA was amplified for 11-13 cycles before being cleaned up using SPRIselect beads. Samples cDNA concentration was determined using a Bioanalyzer. GEX libraries were prepared as recommended by the 10x Genomics Chromium Single Cell 3’ Reagent Kits User Guide (v3.1 Chemistry Dual Index) with appropriate modifications to the PCR cycles based on the calculated cDNA concentration. For sample preparation on the 10x Genomics platform, the Chromium Next GEM Single Cell 3’ Kit v3.1, 16 rxns (PN-1000268), Chromium Next GEM Chip G Single Cell Kit, 48 rxns (PN-1000120), and Dual Index Kit TT Set A, 96 rxns (PN-1000215) were used. The concentration of each library was determined through qPCR utilizing the KAPA library Quantification Kit according to the manufacturer’s protocol (KAPA Biosystems/Roche) to produce cluster counts appropriate for the Illumina NovaSeq6000 instrument. Normalized libraries were sequenced on NovaSeq6000 S4 Flow Cell using the XP workflow and a 50×10×16×150 sequencing recipe according to manufacturer protocol. A median sequencing depth of 50,000 reads/cell was targeted for each Gene Expression Library.

### Mouse Single-cell RNA-sequencing analysis

FASTQ files were aligned to a customized GRCm39 reference genome built with cellranger (v7.0.0) mkref per 10X’s tutorial to include Eyfp and tdTomato (including 3’UTR). Cellranger count with default parameters was employed for sequence alignment and count matrix generation.

Analysis was performed using the Seurat v4.1.1. For quality control, cells with mitochondrial content (percent.mt) exceeding 6% or containing < 3,500 or > 9,500 genes (nFeature_RNA) were removed. After removing unwanted cells and genes, counts were normalized and scaled via the *SCTransform* (51) function. Top 20 principal components (PCs) and resolutions between 0.3 and 0.7 were used for downstream analysis. After preliminary annotation, CAFs were isolated for further analysis using the same workflow. DEGs were evaluated using the Wilcox’s test. GO enrichment analysis was performed using the enrichGO function from clusterProfiler (52) with DEG selected based on the average log2 fold-change weighted by the expression within each cluster (avg_log2_FC *pct.1).

### Human scRNA-seq analysis

Processed data was retrieved from E-MTAB-10607 and GSE176078 (18, 34). Data was processed using Seurat v5.2.1. Cells with percent.mt > 10%, log10 nCount_RNA < 2 or > 5, and log10 nFeature_RNA < 2 or > 4 were removed. After log-normalization and feature selection using mvp, data was scaled by regressing out percent.mt and nCount_RNA. Sample-wise batch-effect was regressed out using harmony. Top 40 PCs and resolutions between 0.1 and 1 were used for downstream analysis. The CAF clusters (PDGRFβ^+^) were isolated and merged from both datasets to identify the PDGFRα^+^PDGRFβ^+^ sub-clusters for further analysis using the same workflow with additional correction for dataset differences. DEG and GO analyses were performed similar to mice data. To identify the human ortholog of mouse Osx^+^myCAF, top 50 DEG and 10 curated markers were used to build the gene set for cell type signature mapping via GSEA using fgsea (53). To create the reference matrix for deconvolution using CIBERSORTx (38), full datasets from both publications were integrated and processed using the same workflow as described above. Annotation of non-CAF clusters was performed over preliminary clusters with annotation of CAF subpopulation transferred from the CAF subset.

### Statistical analysis

All data were analyzed and plotted using GraphPad Prism. Please refer to figure legends for details on statistical tests used for any specific analysis.

## Acknowledgments

We thank the Genome Technology Access Center at the McDonnell Genome Institute at Washington University School of Medicine for help with genomic analysis. The Center is partially supported by NCI Cancer Center Support Grant #P30 CA91842 to the Siteman Cancer Center from the National Center for Research Resources (NCRR), a component of the National Institutes of Health (NIH), and NIH Roadmap for Medical Research.

We thank the Musculoskeletal Histology Core of Musculoskeletal Research Center supported by the National Institutes of Health P30 Grants AR057235 and P30 AR074992, the Washington University Bright Institute, Molecular Imaging Center (NIH P50 CA94056ADD) and Siteman Flow Cytometry Core supported by NCI Cancer Center support grant P30 CA91842 and P30CA091842. This research was supported by grants from National Institutes of Health Grants R01 AR066551 (to R.F.), CA270030 (to R.F.) and CA235096 (to R.F.), grants from Shriners Hospital 85170 and P19-07408 CR (to R.F.), and the Siteman Investment Program, Siteman Cancer Center (Pre-R01 Program to R.F.) and the Foundation for Barnes-Jewish Hospital (3770 and 4642). We thank the St. Louis Breast Tissue Registry at Washington University School of Medicine, Department of Surgery for assistance in obtaining tissue samples. G.F. was supported by Cancer Research Institute postdoctoral fellowship CRI5013, EE was supported by T32GM139774 and F31CA284858, and J.Y. was supported by T32CA113275 and F31CA271721-01. GDL was supported by NIH grant R01 CA254060 and V.M. by F32CA27521. S.A.S. was supported by NIH grants R01 AG059244, CA217208, CA282810. The U.S. Army Medical Research Acquisition Activity, 820 Chandler Street, Fort Detrick, MD 21702-5014, is the awarding and administrating acquisition office, and this was supported in part by the Office of the Assistant Secretary of Defense for Health Affairs, through the Breast Cancer Research Program, under award No. BC181712. Opinions, interpretations, conclusions, and recommendations are those of the authors and are not necessarily endorsed by the Department of Defense. This work was also supported by the Siteman Cancer Center Investment Program (NCI Cancer Center Support Grant P30CA091842), Fashion Footwear Association of New York, and the Alvin J. Siteman Cancer Center Siteman Investment Program (supported by The Foundation for Barnes-Jewish Hospital, Cancer Frontier Fund, to S.A.S.). Schema figures were created in BioRender (Faccio, R. (2025) https://BioRender.com/a54q631).

This publication is solely the responsibility of the authors and does not necessarily represent the official view of NCRR or NIH.

## Extended Data

**Suppl. Fig.1.**
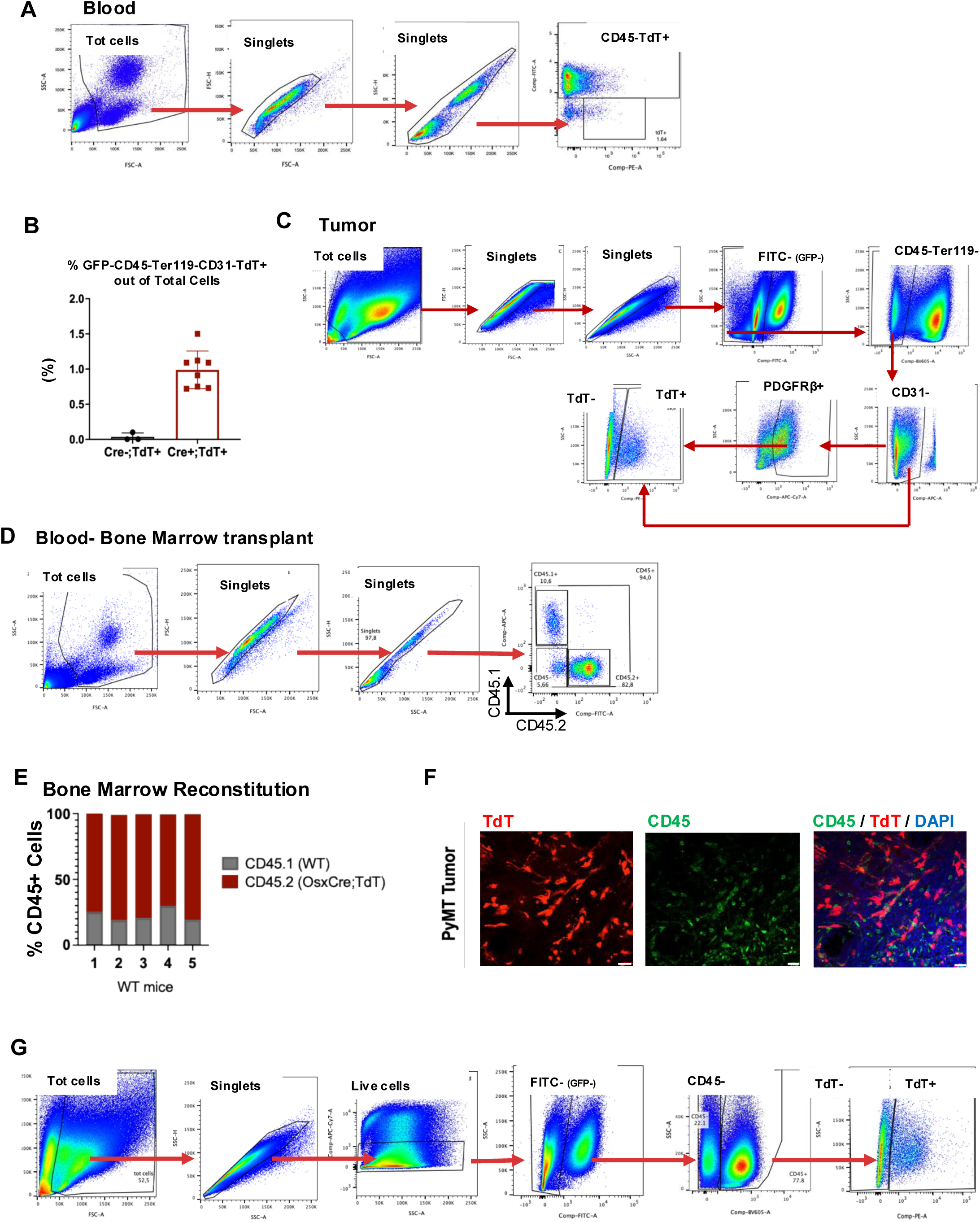
**A**, Gating strategy used for sorting and to determine the percentage of TdT^+^OsteoLin in circulation. **B,** Percentage of TdT^+^OsteoLin in the tumor. **C,** Gating strategy used to determine percentage of CD45^neg^PDGFRβ^+^TdT^+^OsteoLin cells in the tumor. **D-E**, Gating strategies and percentage of chimerism in blood based on CD45.2 and CD45.1 staining. **F,** Immunofluorescence (IF) image of PyMT tumors showing staining for CD45 (green) and DAPI (blue). TdT^+^ cells are displayed in red. **G,** Gating strategy used for sorting tumor-derived CD45^neg^TdT^+^ and CD45^neg^TdT^neg^ for co-injection experiments.

**Suppl. Fig 2.**
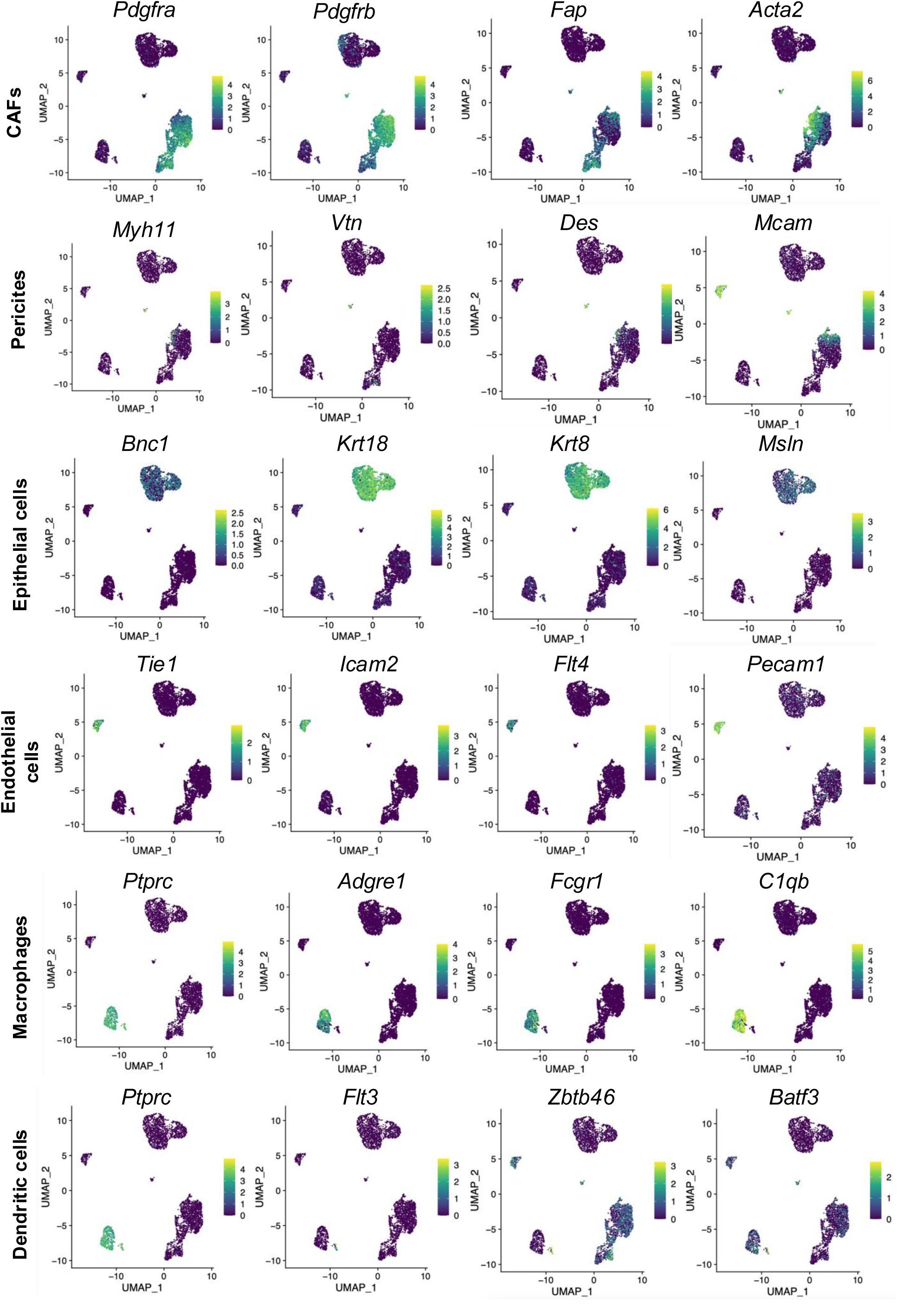
Feature plot of the subpopulation markers.

**Suppl. Fig 3.**
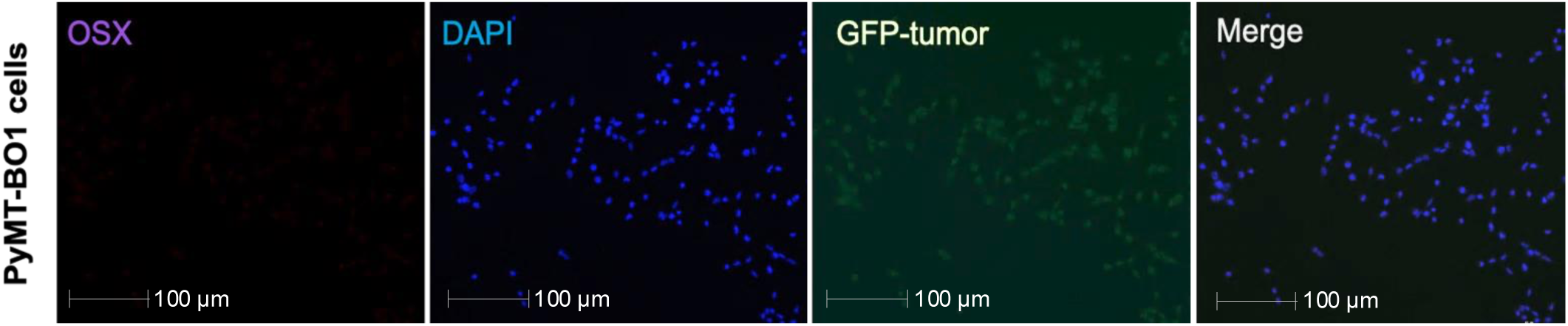
Immunofluorescence was performed on PyMT-BO1-GFP+ cells, stained for Osx (violet) and DAPI (blue).

**Suppl. Fig 4.**
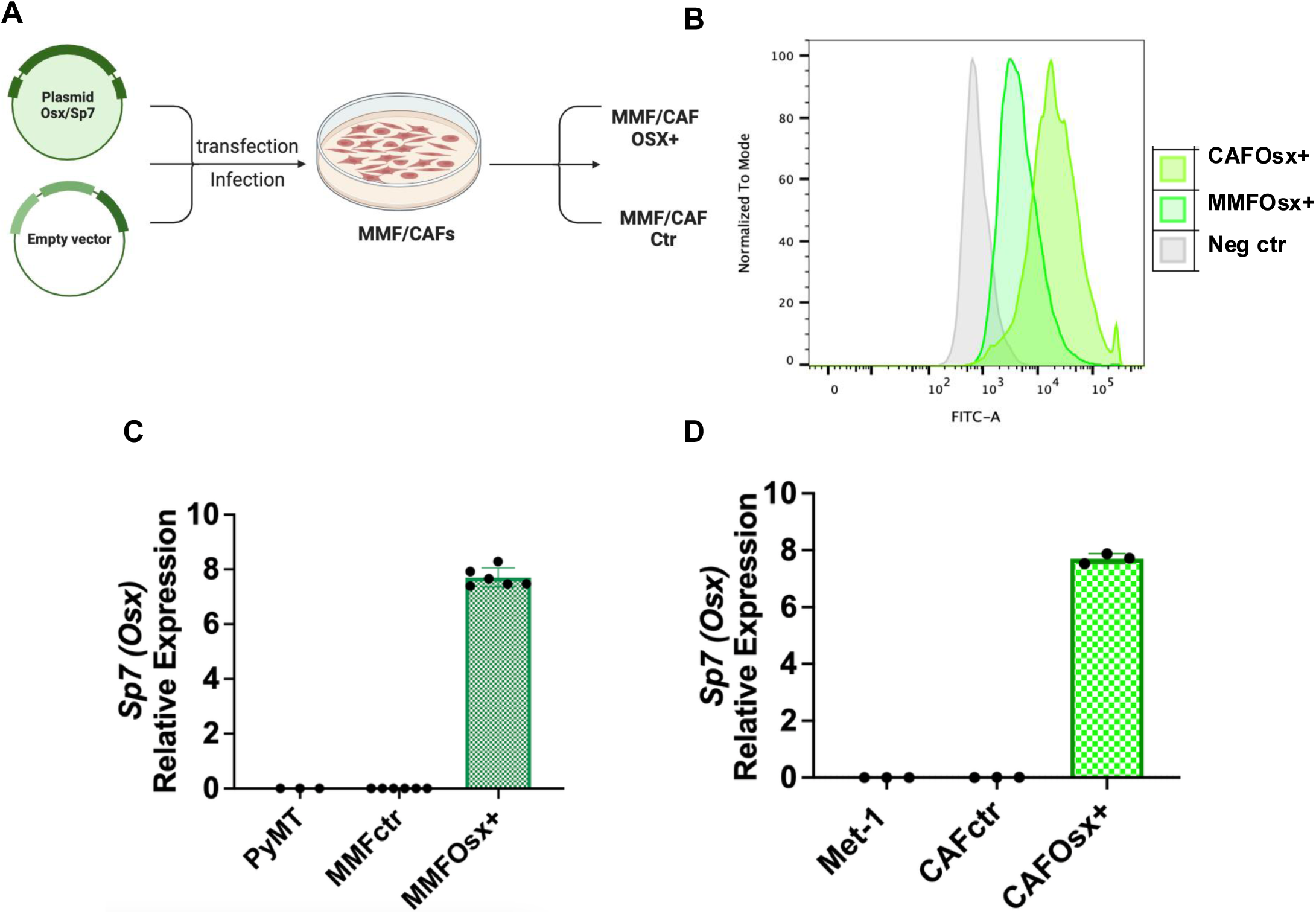
**A,** Schematic representation of GFP-tagged SP7 plasmid or the corresponding empty vector used to generate MMFCtr, MMFOsx^+^, CAFctr and CAFOsx^+^ **B,** Infection and transfection efficiency determined by flow cytometry through detection of GFP expression in MMFsOsx^+^ and CAFsOsx^+^. **C-D,** *Sp7* expression validated by qPCR in MMFOsx^+^ and CAFOsx^+^. Tumor cells, along with MMF and CAF expressing vehicle vector, were used as controls.

**Suppl. Fig 5.**
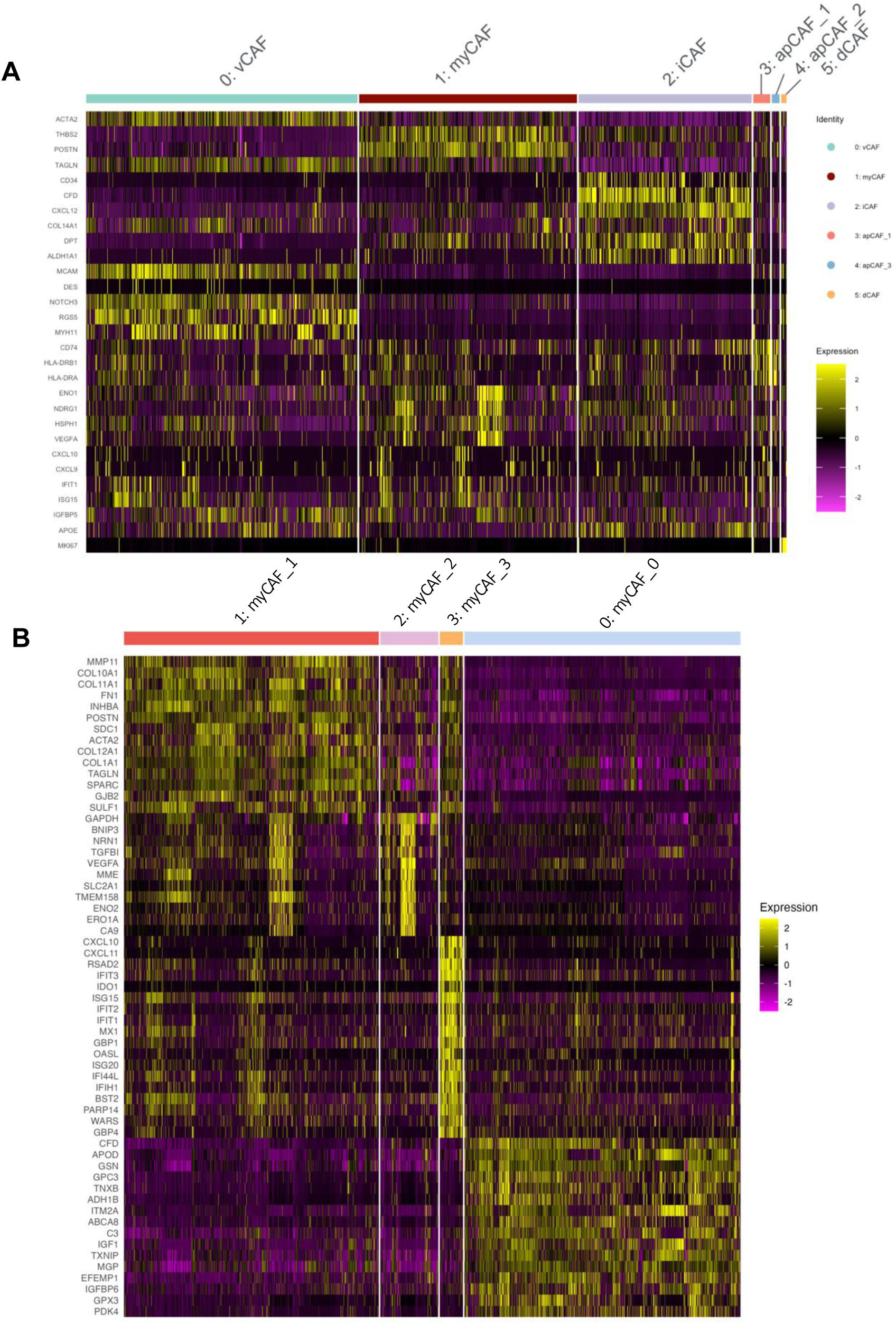
**A-B,** Heatmap showing DEGs among populations.

**Suppl. Fig 6.**
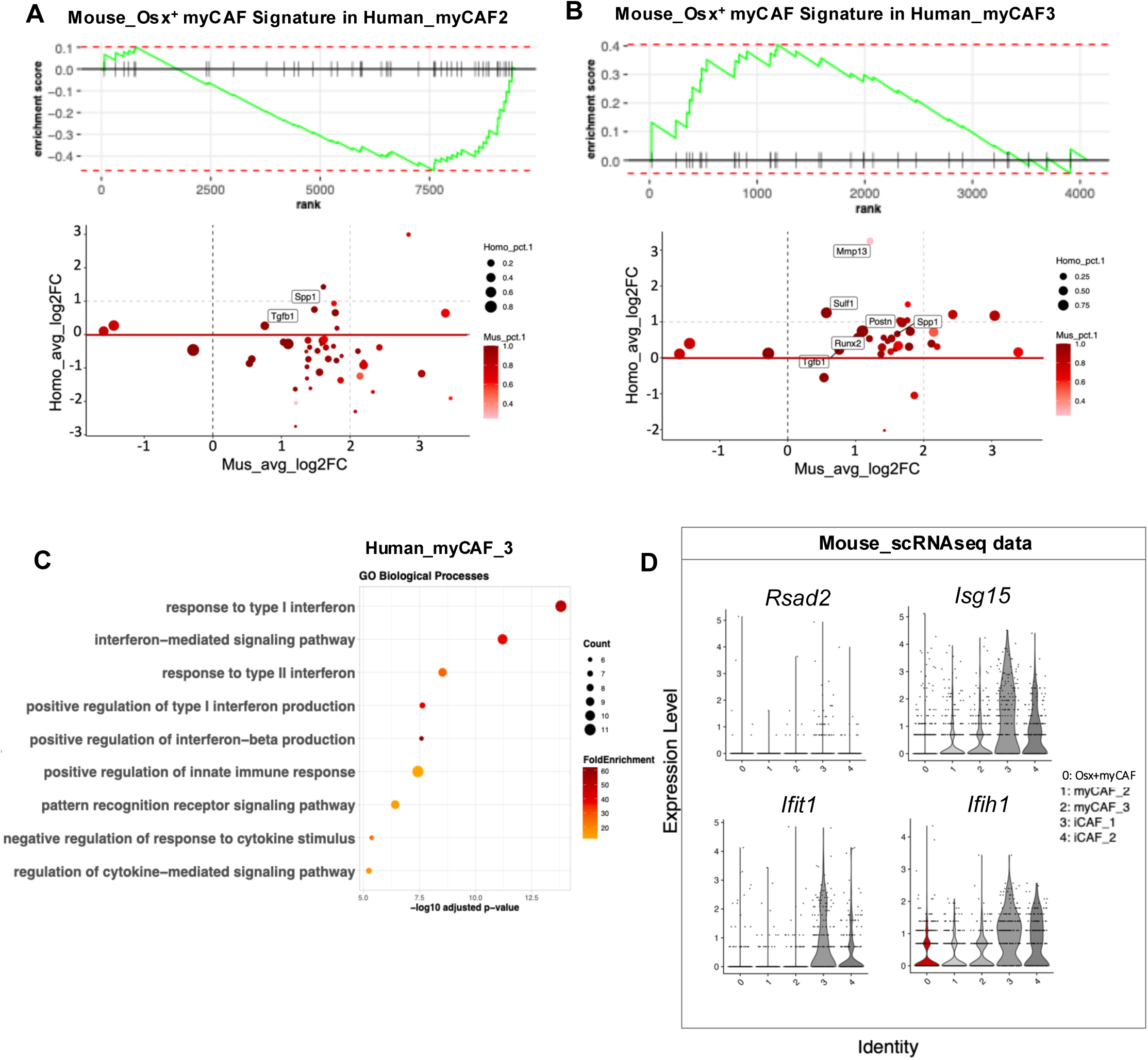
**A-B**, GSEA (top) and scatterplot (bottom) showing the overlap of murine Osx^+^myCAF signature genes with the human myCAF2 (A) and myCAF3 (B) clusters. **C,** Gene Ontology (GO) pathway analyses showig enrichment of genes involved in interferone response in h-myCAF3 population. **D,** Violin Plot showing expression of interferon-related genes in mouse CAF populations.

